# Influenza A virus membrane fusion is regulated by the balance between receptor binding and cleavage

**DOI:** 10.64898/2026.03.25.714262

**Authors:** Steven D. Planitzer, Kevin B. Wu, Zhenyu Li, Mengyang Zou, Poom Ungolan, Na-Chuan Jiang, Balindile B. Motsa, Jia Niu, Tijana Ivanovic

## Abstract

Interactions between influenza A virus (IAV) and its host receptor, sialic acid, influence multiple stages of infection through the opposing activities of hemagglutinin (HA) and neuraminidase (NA). Multivalent HA binding to low-affinity receptors generates high avidity and attachment specificity, while NA-mediated receptor cleavage promotes penetration through sialylated respiratory mucus and progeny virion release. The role of receptor interactions in post-attachment entry, particularly during genome delivery through HA-mediated endosomal membrane fusion, remains unresolved due to conflicting data, limited control of receptor conditions, and difficulty separating attachment from fusion at low efficiency. Here, we address this question using a flow-cytometry based assay that quantifies time-resolved lipid mixing in hundreds of individual virion-membrane pairs per second, combined with programmable control of receptor density and chemistry on target membranes. We show that HA-receptor interactions regulate membrane fusion efficiency by promoting productive fusion-peptide insertion by HA. This effect depends on receptor context: within a defined regime, lipid-mixing efficiency increases with receptor density whereas NA activity reduces it by depleting receptors. Receptor type and HA-receptor binding avidity further modulate lipid-mixing outcomes, establishing a direct link between HA-receptor interactions and fusion efficiency. Together, these results identify receptor binding as an active regulator of membrane fusion and provide a framework that reconciles prior conflicting observations. More broadly, they extend the functional interplay between HA and NA to the level of membrane fusion, with implications for viral adaptation and host specificity.

**Significance Statement:** We resolve a long-standing question by showing that host receptor binding by HA and cleavage by NA regulate influenza A virus membrane fusion, the entry step delivering the viral genome into the cell. HA-receptor engagement promotes productive HA membrane insertion, leading to fusion. Receptor density, HA-receptor affinity, and NA catalytic activity tune fusion outcomes. These results were enabled by two methodological advances: precise control of receptor presentation on target membranes and sensitive measurement of attachment and membrane fusion at the single-virion level. By defining how the opposing activities of HA and NA regulate fusion, this study extends their functional interplay to a key step in entry and provides new insight into how these proteins coevolve during viral adaptation.

## Introduction

Interactions between influenza A virus (IAV) and its host receptor, sialic acid, influence multiple stages of infection through the opposing actions of two virion surface glycoproteins. Hemagglutinin (HA) binds receptors whereas neuraminidase (NA) cleaves them, and their activities are evolutionarily constrained to remain balanced (1, 2). HA mediates virion attachment to cells leading to endocytosis while NA promotes penetration through sialylated respiratory mucus and progeny virion release after budding (3–7). The HA-NA functional balance is constrained by the distribution of sialic-acid species in each host or tissue within a host. Bird and human cells are enriched in α2,3- and α2,6-linked sialic acids, respectively, and dominant receptor types vary across tissues and cell types in the airway (8, 9). Strain-dependent differences in the binding and cleavage of these receptor types thus contribute to host and tissue tropism (9–14), and changes in these relative activities can facilitate cross-species transmission of pandemic potential.

Beyond its role in attachment, HA facilitates membrane fusion in endosomes. The fusion competent form is a trimer of disulfide-linked HA1-HA2 heterodimers generated by proteolytic cleavage of the inactive HA0 precursor (15). Both receptor binding, mediated by HA1, and membrane fusion, mediated by HA2, depend on high HA density on the virion surface (16). Individual HA-receptor binding affinities are low (*k*_d_ ∼3mM), and attachment depends on multivalent interactions (17, 18). Fusion requires 3-4 neighboring HA2 molecules to engage the target membrane. Upon endosomal acidification, individual HAs undergo stochastic conformational activation in which reversible HA1 dilation is tightly coupled to the probability of HA2 extension (Fig. 1*A*) (19–22). The N-terminal fusion peptides of HA2 are released and projected beyond the HA1 domain, inserting into the target membrane and forming a long-lived extended intermediate (Fig. 1*A*, *right*). This step is rate-limiting (22). Subsequent refolding to the energetically favorable post-fusion conformation (‘foldback’) occurs rapidly and drives membrane fusion (22). Importantly, the hydration-force barrier is overcome only when 3-4 neighboring HA trimers insert and undergo coordinated foldback (Fig.1*B*) (22–24).

**Figure 1.**
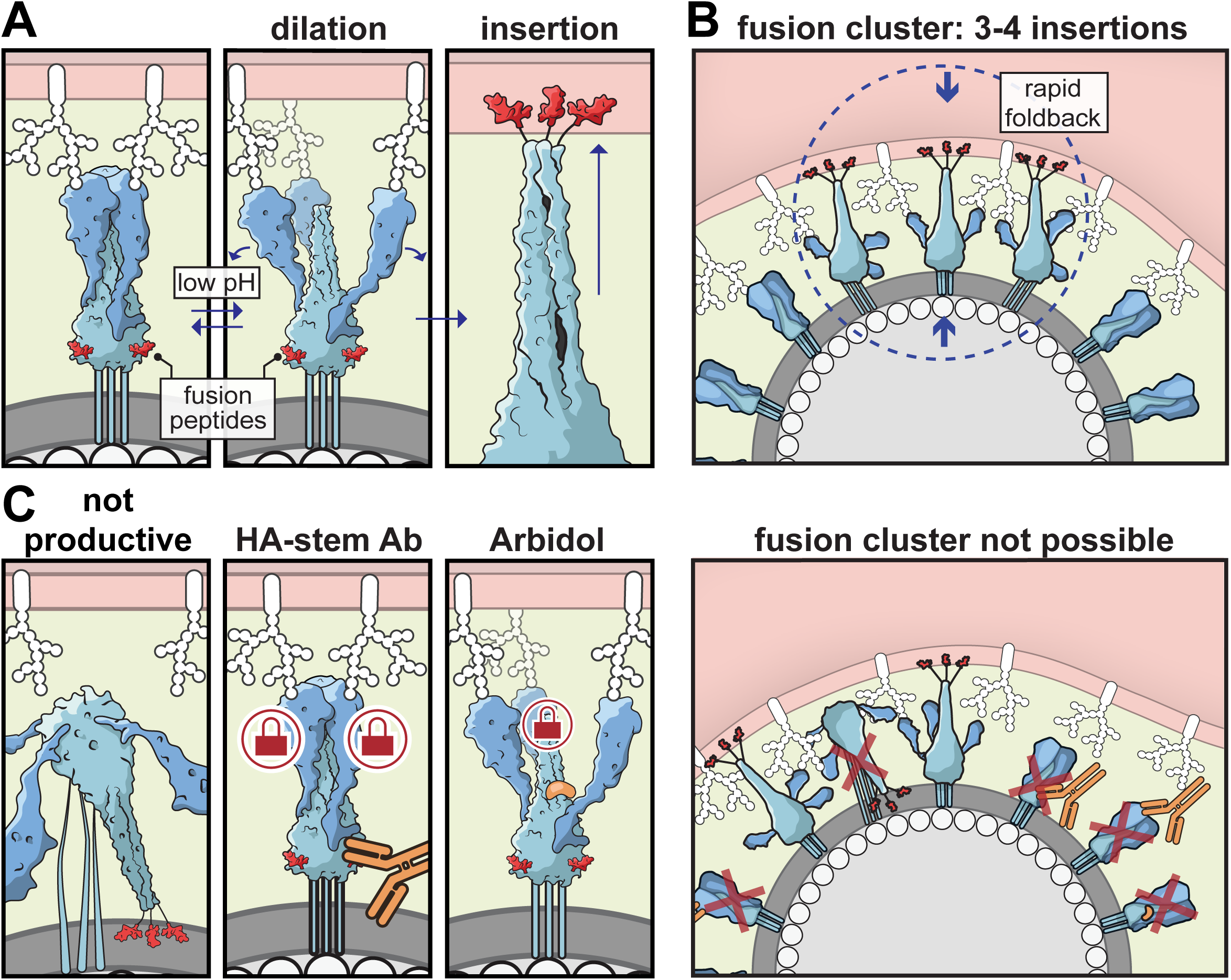
The mechanism of HA-mediated membrane fusion. (*A*) Endosomal acidification induces reversible HA1 opening, increasing the probability of fusion-peptide release (*left* and *middle*). This release is rate limiting, and lower pH prolongs the HA1 open-state dwell time. Fusion-peptide release and membrane insertion generate a long-lived extended intermediate, as single inserted HAs cannot complete foldback (*right*). (*B*) Overcoming the hydration-force barrier requires insertion and coordinated foldback of 3-4 neighboring HA trimers (a fusion cluster). Insertions occur stochastically within the virion-target contact patch. Once the critical number of neighboring HAs has inserted, foldback proceeds rapidly. (*C*) HA inactivation constrains the geometries available for fusion-cluster assembly. Some HAs undergo nonproductive refolding without target-membrane insertion, stem-binding antibodies stabilize HA in the prefusion conformation, and Arbidol delays HA2 extension beyond the transient window of HA1 opening (*left*). When enough HAs are rendered unavailable, fusion-cluster formation becomes impossible and fusion is inhibited (*right*). Some graphical elements are adapted from (60).

The virion-target contact patch contains tens to thousands of HAs depending on virion size, but only a fraction of HAs can participate in fusion (25). A patch might include inactive HA0 or NA in place of active HA, or some active HAs might transition to the post-fusion state without inserting (Fig. 1*C*) (26); high HA density buffers these intrinsic inefficiencies. Fusion is inhibited when fusion clusters cannot assemble, such as under additional inhibitory pressure (26) (Fig. 1*C*). For example, stem-binding HA antibodies lock HA in the prefusion conformation (27, 28) whereas Arbidol, a broad-spectrum fusion inhibitor, delays HA2 extension beyond the transient window defined by HA1 dilation in endosomes (20) (Fig. 1*C*). The potential for additional regulatory events within the contact patch, such as receptor binding and cleaving, is intriguing but remains largely unexplored.

The role of sialic acid receptors in IAV membrane fusion has been debated for decades. Some studies concluded that receptor-bound HAs cannot directly participate but might indirectly promote fusion by facilitating HA clustering in the contact patch (29, 30). Others proposed that sialic acids play no role in virus entry post attachment, reporting no change in fusion kinetics for virions recruited to membranes nonspecifically or via poorly matched sialic-acid moieties (31–33). In contrast, other studies reported notable though varied effects of HA-receptor interactions on fusion, such as by influencing fusion kinetics (34, 35), altering HA conformational dynamics (36), facilitating fusion-peptide insertion in general (37) or in crowded membrane environments (38), enhancing fusion efficiency (39), or fundamentally restructuring the fusion pathway (35). The varied and sometimes inconsistent findings of prior studies suggest that receptor effects on membrane fusion are complex and might be sensitive to context. Beyond biological variability, evaluating the role of glycan receptors in membrane fusion poses distinct technical challenges. Tools to precisely vary glycan-receptor type or density in fusion assays are lacking and low receptor densities or suboptimal receptors reduce virion binding, limiting reliable quantification. Moreover, bulk measurements cannot resolve receptor effects on fusion independently of attachment. This combination of biological complexity and technical challenge underscores the need for tools that enable precise control of the receptor environment and high-statistical-power quantification of rare binding and fusion events at single-virion resolution.

We developed a flow cytometry-based assay that quantifies time-resolved distributions of lipid-mixing states among single virions bound to individual cells or vesicles and enables measurements of lipid mixing under low-attachment and low-efficiency regimes. To control the receptor environment during fusion, we engineered programmable DNA oligonucleotide-mediated displays of synthetic receptors on target membranes. We demonstrate that receptor density and type are key determinants of membrane fusion efficiency. Fusion is favored by high HA-receptor affinity and high receptor density and disfavored by NA activity. Our results support a model in which receptor binding favors HA insertion and reduces the incidence of spontaneous inactivation. In sum, our results indicate that receptor binding is not merely a passive determinant of virion attachment but also a regulator of membrane fusion efficiency and suggest a framework to reconcile previously conflicting observations.

## Results

### IAV membrane fusion is sensitive to receptor availability within the contact patch

While both attachment and membrane fusion require high HA density on virions, only attachment is known to depend directly on multivalent receptor engagement. Whether fusion remains sensitive to receptor availability within the contact patch after attachment remains unclear. We therefore developed a novel approach, Single-Pair Cytometric Analysis of Lipid Mixing (spCALM), that quantifies fusion independently of attachment efficiency and is sensitive in low-attachment and low-fusion regimes. spCALM enables tracking of real-time distributional dynamics of bound virion-membrane target pairs undergoing lipid mixing in suspension (Figs. 2*A*,*B*, S1, and Movie 1). Here, cell or vesicle targets are incubated with R18-labeled virions at a ratio of 0.15 virions per target to allow attachment of at most a single virion. In the analysis, free targets are distinguishable from virion-bound ones based on R18 fluorescence and binding is quantified as the percentage of targets with bound virions (Fig. 2*B*). Fusion is triggered by dilution into a low-pH buffer shortly after initiation of flow, and lipid mixing is detected as the emergence of a brighter, R18-dequenched population. Lipid-mixing efficiency is quantified as the fraction of lipid-mixed pairs at the reaction plateau (Figs. *2A-B*, S2). As an initial validation, spCALM with erythrocyte ghosts and H3N2 IAV at pH 5.54 yielded complete (∼100%) lipid mixing (Fig. S2*A*). In contrast, H1N1 IAV was less efficient, exhibiting ∼3% and ∼33% lipid-mixing efficiency at pH 5.69 and 5.33, respectively (Fig. S2*B*). These results are consistent with prior fusion studies on supported planar membranes showing lower baseline efficiency of H1N1 IAV and a higher stoichiometric requirement in the fusion cluster – 3 neighboring HAs for H3N2 versus 4-5 for H1N1 (26, 27).

**Figure 2.**
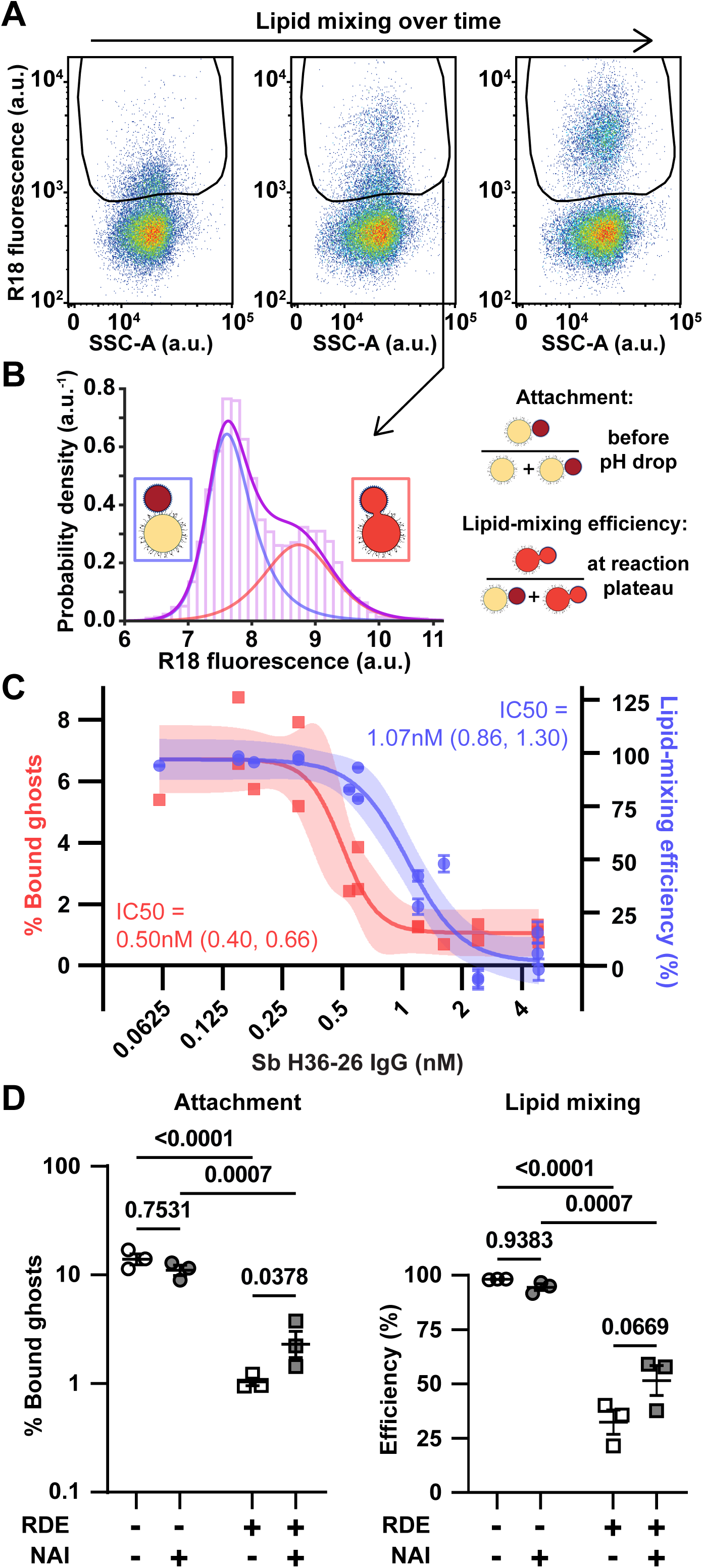
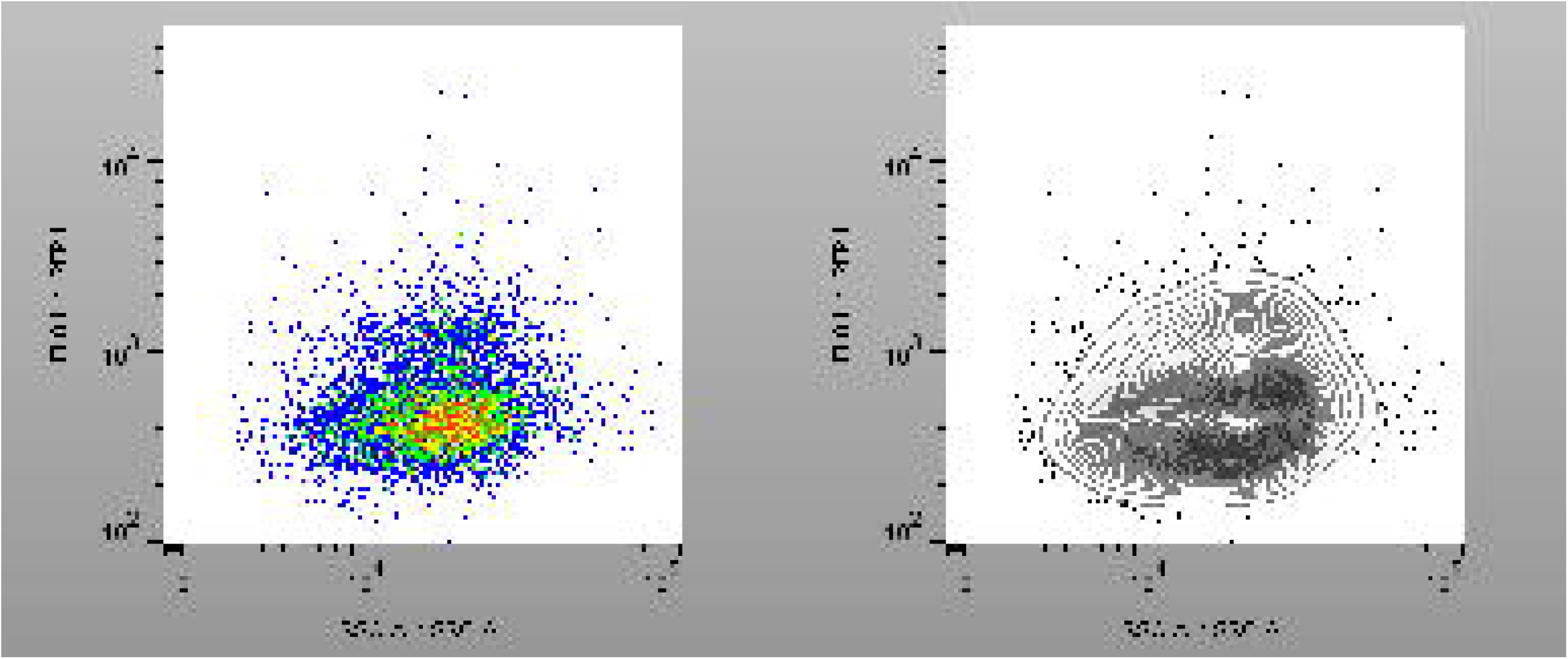
spCALM with erythrocyte ghosts reveals membrane fusion is inhibited by an HA1-binding antibody and reduced receptor density. (*A*) Example spCALM pseudocolor scatter plots of R18 fluorescence (*y-axis)* versus blue-laser side scatter (*x-axis*). Event distributions from three acquisition windows reveal characteristic subpopulations of R18-labeled H3N2 virions with erythrocyte ghosts before pH drop (*left*), early after acidification (*middle*), and at the reaction plateau (*right*) (see Movie 1). The gate identifies bound and lipid-mixed virion-ghost pairs. Lipid mixing between virions and ghosts dequenches R18 fluorescence, shifting lipid-mixed pairs to a brighter population distinct from unfused pairs. Acquisition intervals were selected to display a comparable number of events. (*B*) Analysis of spCALM data. (*Left*) The histogram shows the fluorescence distribution of virion-ghost pairs at an intermediate timepoint (data from *top row, middle panel*). Unfused (*blue line)* and lipid-mixed (*red line*) reference distributions were obtained by Burr fits to the initial and final states; their relative weights at each time bin quantify lipid mixing to derive the reaction kinetics. (*Right*) Attachment is expressed as the percentage of cells with bound virions before pH drop. Lipid-mixing efficiency is defined as the fraction of lipid-mixed pairs among total pairs at the reaction plateau. (*C*) Differential inhibition of attachment and lipid mixing by HA1-binding antibody H36-26. H1N1 virions were pretreated with increasing antibody concentrations, and binding and lipid mixing were quantified by spCALM in the continued presence of antibody. The 0 nM condition was included in curve fitting but not plotted. Data points deriving from three independent experiments were separately plotted. Error bars (lipid mixing) indicate measurement uncertainty. Curves are logistic fits to the combined data with shaded 95% confidence intervals. (*D*) Inhibition of attachment and lipid mixing by reduced receptor density. Attachment (*left*) and lipid mixing (*right*) of H3N2 virions with erythrocyte ghosts at 34°C and pH 5.54 in the presence or absence of 100 nM NA-inhibitor oseltamivir (NAI), measured by spCALM. Ghosts were treated with RDE to reduce receptor density or mock-treated. Plots show geometric mean +/- SEM (*attachment*) and arithmetic mean +/- SEM (*lipid mixing*) with individual data points from three independent experiments. Statistical significance was assessed by Tukey’s multiple-comparisons test.

We next used spCALM to test whether attachment-blocking antibodies inhibit membrane fusion of virions that have successfully attached. We measured attachment and lipid mixing of H1N1 IAV virions with erythrocyte ghosts in the presence of increasing concentrations of H36-26 IgG, a strongly neutralizing HA1-binding antibody. H36-26 dose-dependently inhibited not only the attachment but also the lipid-mixing efficiency of pre-bound pairs (Fig. 2*C*). More antibody was required to reach half-maximal inhibition for lipid mixing than for attachment. Lipid-mixing lag time, defined as the time to median lipid mixing, also increased in the presence of H36-26 IgG, although these measurements were highly variable (Fig. S3). Inhibition of lipid mixing by bound virions might result from reduced virion-receptor interactions within the contact patch or from steric hindrance by antibodies that prevents HA2 fusion peptides from reaching the ghost membrane. In either case, these results indicate that head-binding antibodies can inhibit membrane fusion independently of their effects on attachment.

To directly test whether membrane fusion is sensitive to virion-receptor interactions, we performed spCALM experiments using erythrocyte ghosts pretreated with *Vibrio cholerae* receptor-destroying enzyme (RDE). Reduced receptor density was confirmed by decreased binding of fluorescently labelled α2,6-specific SNA-lectin (Fig. S4*A*). We quantified attachment and lipid-mixing efficiency of H3N2 IAV virions on RDE- or mock-treated ghosts in the presence or absence of 100nM oseltamivir carboxylate, a small-molecule NA-inhibitor (NAI). RDE treatment markedly reduced attachment and lipid-mixing efficiency (Fig. 2*D*). Lag time increased, but this effect was more variable (Fig. S4*B*). NAI had no effect under receptor-rich conditions (mock-treated ghosts) (Fig. 2*D* and S4*B*). In contrast, under receptor-depleted conditions (RDE-treated ghosts), NAI modestly increased attachment and lipid mixing relative to vehicle control (p = 0.038 and 0.067, respectively) but did not affect lag time. Together, these experiments demonstrate that membrane fusion is sensitive to receptor density within the contact patch but do not directly link specific protein-receptor interactions to lipid mixing.

### Programmable glycan-receptor displays for membrane-fusion studies

To dissect receptor-dependent molecular events within the contact patch, we developed a programmable receptor-display platform in which receptor density, type, and distance from the target membrane can be precisely controlled. We used double-stranded DNA (dsDNA) as a defined-length spacer between the target membrane and receptors. Its rigidity (persistence length of tens of nanometers), and sequence-independent axial rise (∼0.34 nm per base pair in B-form) allow dsDNA to function as a molecular ruler of specified length (40). Prior work demonstrated that lipid-anchored dsDNA tethers can link viral and target membranes while preserving fusion (31), further motivating our use of dsDNA spacers to display receptor mimetics in a programmable geometry. Native IAV receptors are glycoproteins or glycolipids containing polysaccharide structures with a terminal *N*-acetylneuraminic acid (Neu5Ac, generally known as sialic acid) and a penultimate galactose with either an α2,3-glycosidic linkage (avian) or an α2,6-glycosidic linkage (mammalian) (41, 42). To mimic the native receptor, we chose a commercially available trisaccharide sialyllactose consisting of terminal sialic acid linked to galactose by either an α2,3-glycosidic bond (3’SL) or an α2,6-glycosidic bond (6’SL). Our receptor mimetic is comprised of two separate components: a lipid-ssDNA for membrane anchoring and a sialyllactose-conjugated complementary ssDNA strand for virion engagement (receptor-ssDNA), allowing for programmable modulation of receptor type with full control over the spacer length (Fig 3*A*). This platform allows precise tuning of receptor density by substituting receptor-conjugated ssDNA with receptor-free ssDNA at defined ratios (Fig. 3*B*).

**Figure 3.**
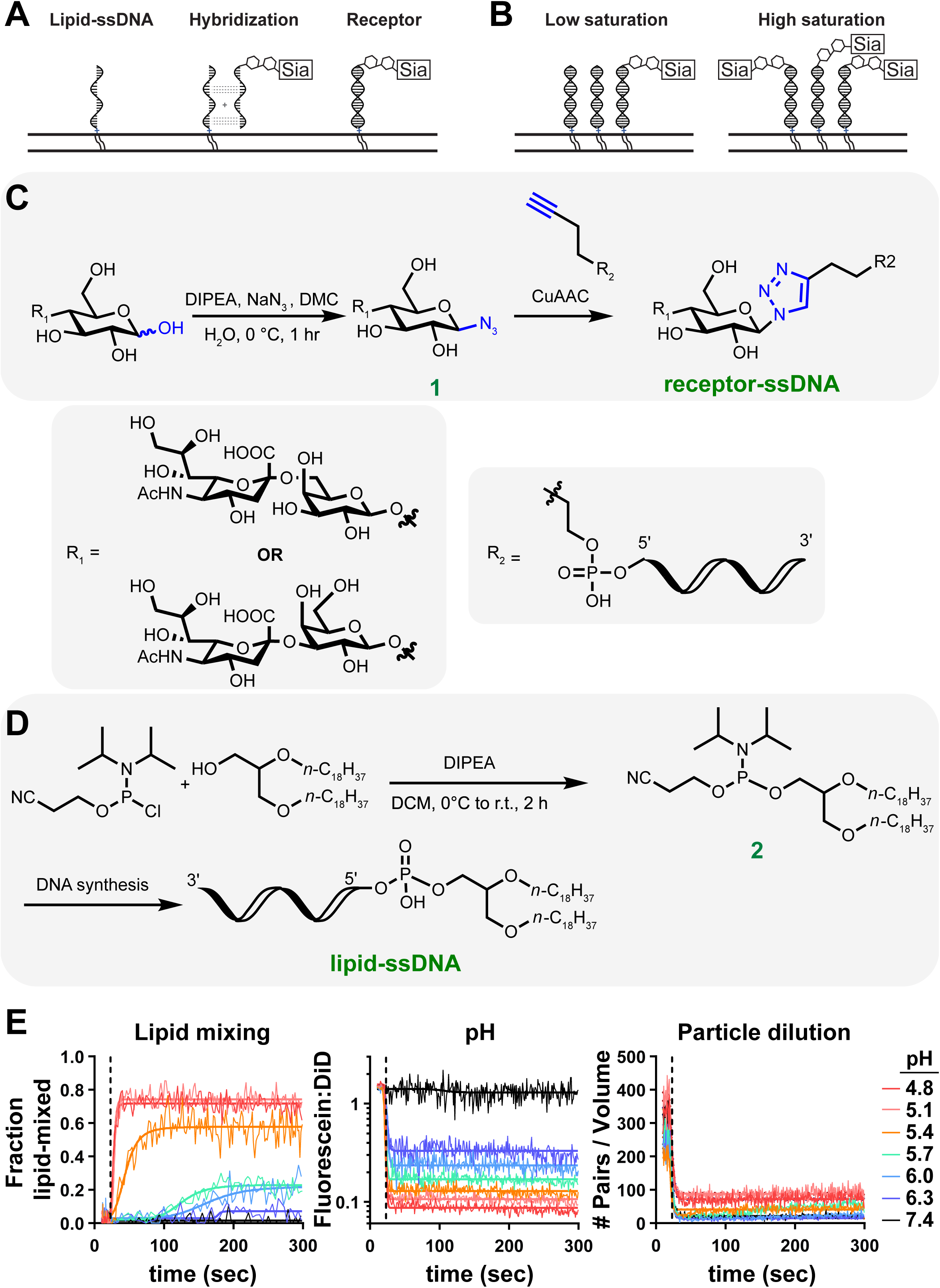
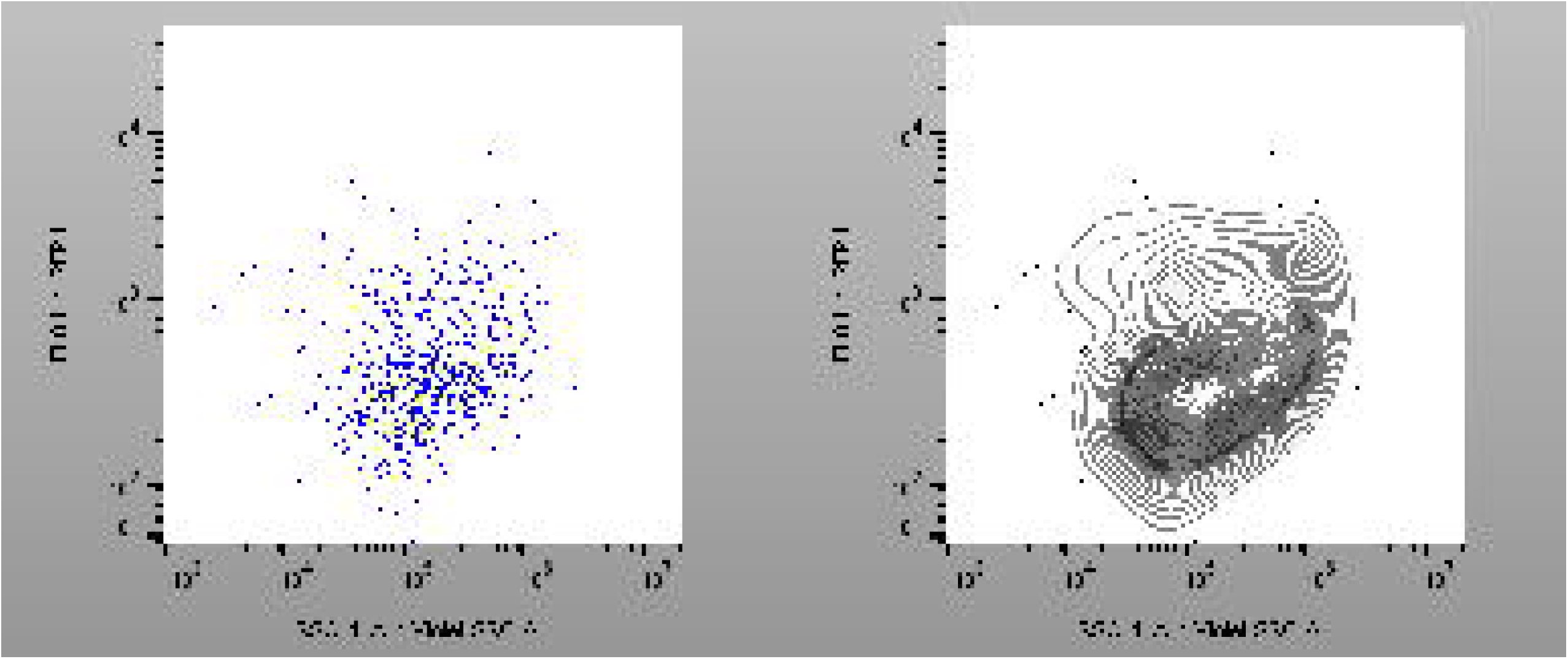
Programmable receptor display on synthetic membranes and its validation by spCALM. (*A*) Lipid-ssDNA is incorporated into liposomes at 1 mol% and hybridized to receptor-ssDNA. (*B*) Receptor density is controlled by changing the ratio of receptor-ssDNA to free ssDNA. Complete list of ssDNA sequences is included in Supplementary Table 1. (*C*) Synthesis of receptor-ssDNA. Azido-sialyllactose (**1**) was generated by azidation of commercial sialyllactose with either an α2,3 or α2,6 glycosidic bond. copper-catalyzed Huisgen azide–alkyne cycloaddition (CuAAC) of **1** with 5’-hexynyl-modified ssDNA of the desired nucleotide spacer length then yielded ssDNA-conjugated receptor. (*D*) Synthesis of lipid-ssDNA. Lipid phosphoramidite (**2**) was generated by reacting 1,2-*O*-dioctadecyl-*rac*-glycerol with 2-cyanoethyl *N,N*-diisopropylchlorophosphoramidite. A DNA-synthesizer then conjugated the desired oligonucleotide sequence, with **2** serving as the last “base”. (*E*) Real-time spCALM traces for H3N2 virions and 6’SL-receptor liposomes across a range of pH values. Traces begin at dilution into buffers of the indicated pH. The dotted vertical line indicates the measurement lag. (*Left*) Fraction of lipid-mixed pairs as a function of time. Lag time is defined as the time to the median lipid-mixing extent. (*Middle*) Sample acidification is reflected in the ratio of median fluorescence of the pH-sensitive dye (fluorescein) to the pH-stable membrane dye (DiD) incorporated in liposomes. (*Right*) Dilution of virion-liposome pairs identifies the measurement lag.

To generate receptor-ssDNAs, the reducing end of a commercially available 3’- or 6’-sialyllactose (SL) was azidated using sodium azide in the presence of 2-chloro-1,3-dimethylimidazolinium chloride (DMC) (43). The resulting azidosialyllactose **1** was then conjugated to alkyne-functionalized ssDNA via copper-catalyzed Huisgen azide-alkyne cycloaddition (CuAAC) reaction (Fig. 3*C*). The lipid-ssDNAs were constructed following standard automated DNA synthesis procedures with a lipid phosphoramidite (1,2-*O*-dioctadecyl-rac-glycerol-2-cyanoethyl-*N*,*N*-diisopropylchlorophosphoramidite) **2** coupled to the 5’-end of the ssDNA sequence (Fig. 3*D*). Receptor-ssDNA and lipid-ssDNA were characterized by MALDI-TOF mass spectrometry and LC-MS mass spectrometry, respectively, and the observed mass was consistent with the expected values for these compounds (Fig. S5).

For initial validation, we implemented the receptor displays in a Total Internal Reflection Fluorescence microscopy (TIRFm)-based supported planar bilayer (SPB) assay (20, 22, 24, 25) (Fig. S6). Briefly, we incorporated lipid-ssDNA into SPBs at 1mol% and hybridized them to complementary 6’SL-ssDNA (full, or 100% saturation), receptor-free ssDNA (0% saturation), or a 1:6.25 mixture of 6’SL-ssDNA and receptor-free ssDNA (16% saturation). Receptor-rich SPBs efficiently recruited virions and attachment was specific (Fig. S6, compare 100% and 0% receptor saturation). However, at lower receptor densities (16% receptor saturation), too few virions attached rendering SPB-based single-virion imaging impractical for quantitative lipid-mixing analysis.

To circumvent this limitation, we next employed spCALM with liposomes in suspension. In addition to the receptor displays, liposomes also incorporated pH-stable and pH-sensitive fluorescent dyes to facilitate population gating and monitor acidification (Fig. S7). We validated spCALM with liposome targets by testing H3N2 IAV across a range of pH values at 60% receptor saturation. As expected, decreasing pH accelerated lipid mixing and increased the plateau efficiency (Fig. 3*E*). After a brief delay due to instrument dead-volume (measurement lag), the diluted reaction reached the detector and lipid mixing was monitored in real time (Fig. 3*E*, *left*). At physiologically relevant pH (≥pH 5.4), lipid mixing was slow relative to this delay, permitting derivation of average lag times and, at pH 5.7 and 6, reliable model fitting and comparison with prior single-virion SPB fusion studies (Fig. S8). In agreement with SPB-based experiments (22, 24, 26, 27), lipid-mixing kinetics were well fit by cumulative gamma probability density functions with 3-4 underlying steps. Importantly, spCALM enabled measurement of low-efficiency lipid mixing that is otherwise inaccessible to SPB-based techniques. We further tested 12- and 24-nucleotide (nt) dsDNA-spacer lengths and found no difference in lipid-mixing efficiency or kinetics, although binding was more efficient with the 24-nt spacer (Fig. S9). We used 24-nt spacers to display receptors in the remaining experiments in this study. The combined experiments validate our receptor-display platform in combination with spCALM for extracting lipid-mixing kinetics and efficiency. spCALM preserves the single-virion resolution of TIRFm-based approaches and makes possible rapid evaluation of distributional dynamics with improved throughput, statistical power, and sensitivity to rare events.

### NA-inhibition promotes lipid mixing under receptor-limited conditions

Our spCALM data with erythrocyte ghosts suggested that NA-mediated receptor cleavage might attenuate membrane fusion under receptor-limited conditions (see Fig. 2*D*). We next applied our programmable receptor-display platform to probe the effect of NA catalytic activity under defined receptor conditions. We measured lipid mixing by H3N2 IAV over a range of NAI concentrations with liposomes incorporating low receptor density (5% saturation) or only receptor-free DNA. We detected no lipid mixing for liposomes without receptors (Fig 4*A*), confirming that neither low level of nonspecifically bound virions (Fig. S10) nor unbound virions contribute to the lipid-mixing signal under our assay conditions. For liposomes with receptors, NAI did not significantly alter virion attachment or lipid-mixing lag time (Fig. S10), but it enhanced lipid-mixing efficiency in a dose-dependent manner with EC50 of 0.015 nM (0.003, 0.130) (Fig 4*A*). This EC50 is comparable to the NAI IC50 for this virus in the MUNANA assay (∼0.09 nM), which measures virion-associated NA activity via fluorogenic substrate cleavage (Fig. S11). These results are consistent with NAI promoting lipid mixing by inhibiting viral NA-mediated receptor cleavage on target liposomes. The combined data reveal that viral NA activity antagonizes fusion under receptor-limited conditions and further implicate receptor density as a determinant of membrane fusion.

**Figure 4.**
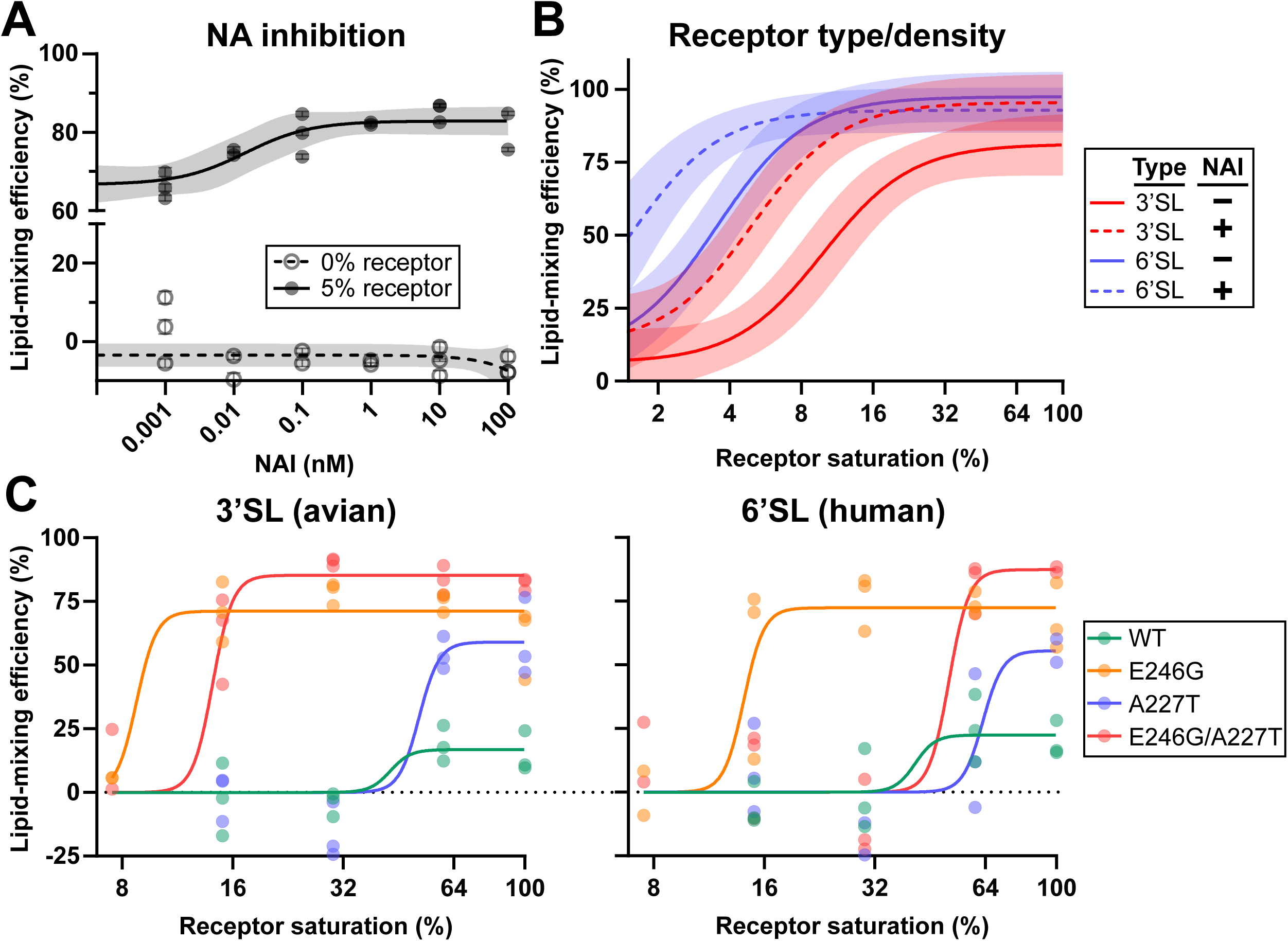
Receptor density, type, and binding avidity are determinants of lipid-mixing efficiency. (*A*) NAI-dependent promotion of lipid mixing. Liposomes displaying either 6’SL receptors (5% saturation) or no receptors (0% saturation) were incubated with H3N2 virions in the presence of the indicated concentrations of NAI. Fusion was triggered by dilution into pH 5.54 buffer, with NAI present throughout. Data points derive from three independent experiments. Logistic (*5% saturation)* and linear *(0% saturation)* curves were fit to the combined data, with shaded 95% confidence intervals. (*B*) Dependence of lipid-mixing efficiency on receptor density, receptor type, and NAI. Lipid mixing of H3N2 virions with liposomes displaying 3’SL or 6’SL receptors at the indicated range of saturations in the presence of absence of 100 nM NAI. Fusion was triggered by dilution into pH 5.68 buffer. Data points derive from three independent experiments. Logistic curves with a shared Hill slope were fit to the combined data from each condition, with shaded 95% confidence intervals. (*C*) Lipid-mixing efficiency of H1N1 WT and HA1 receptor-avidity mutants at pH 5.30. Efficiency was plotted as a function of receptor density for 3’SL receptors (*left*) or 6’SL receptors (*right*). Data points derive from three independent experiments. Logistic curves with a shared Hill slope were fit to the combined data for each virus variant. Efficiency values were offset so that the bottom plateau of each fitted curve is set at 0 to account for differences in baseline signal between the viruses.

### Receptor density and type are determinants of IAV membrane fusion

To directly evaluate the effect of receptor density on membrane fusion, we performed spCALM with H3N2 IAV virions and liposomes displaying a range of 6’SL-receptor saturations in the presence or absence of 100 nM NAI. Lipid-mixing efficiency exhibited sigmoidal dependence on receptor density when plotted on log_2_ scale, having strong dependence over about a four-fold range of receptor densities and reaching plateau values well below full receptor saturation (Fig. 4*B*). Virion attachment was also sensitive to receptor density but over a broader range of values than lipid mixing (Fig. S12, *right*), indicating distinct receptor-density dependencies for the two processes. NA inhibition did not affect lipid-mixing lag time (Fig. S12, *left*) but increased lipid-mixing efficiency and shifted the receptor-density response curve toward lower densities (Fig. 4*B*). Lipid mixing of bound virions is therefore sensitive to receptor density within the contact patch, and protection of limited receptors by inhibiting viral NA activity enhances lipid mixing.

The combined data identify contributions of receptor density and receptor cleavage by NA in influencing lipid mixing within the contact patch but do not directly implicate HA. To begin probing HA’s contribution to receptor-dependent events, we additionally measured lipid mixing of H3N2 IAV with 3’SL receptor-displaying liposomes. Binding avidity for sialic-acid linkages varies by strain, and the H3N2 strain used here preferentially engages α2,6-linked receptors (44) (11). Lipid mixing with 3’SL receptor-displaying liposomes required higher receptor density not only for attachment (Fig. S12, *right*) but also to achieve comparable lipid-mixing efficiency (Fig. 4*B*). As with 6’SL receptors, NA inhibition promoted lipid mixing and shifted the receptor-density response curve toward lower densities. Notably, 6’SL receptors supported lipid mixing at lower densities than 3’SL receptors even under complete NA inhibition (see Fig. S11), suggesting that HA-receptor avidity within the contact patch influences lipid-mixing efficiency (Fig. 4*B*). However, these data do not exclude the possibility that NA might continue to bind receptors in the presence of NAI and thereby contribute to lipid-mixing efficiency independent of its catalytic activity.

### HA-receptor avidity is a determinant of IAV membrane fusion

To directly test whether HA-receptor interactions promote lipid mixing, we generated H1N1 IAV variants incorporating wild-type (WT) NA and HA1 point mutations reported to alter sialic acid-binding avidity (E246G and A227T), either individually or in combination (45). The E246G mutation increases receptor avidity, and A227T in the double mutant restores WT avidity as measured by virion-mediated agglutination of red blood cells with reduced receptors (45). Mutant virus plaque sizes inversely correlated with the expected receptor-binding avidities, likely reflecting avidity-dependent retention of progeny virions on producer cells (Fig. S13*A*). Because these mutations reside near the HA1 receptor-binding site and away from HA2 fusion machinery, we reasoned that differences in receptor-density dependence of lipid mixing for the mutants would likely reflect differences in HA-receptor interactions within the contact patch. We measured attachment and lipid mixing by spCALM using liposomes displaying a range of 3’SL or 6’SL receptor densities in the presence of 100 nM NAI to prevent NA-mediated receptor cleavage (Figs. 4*C* and S13*B*). As with H3N2 IAV, lipid mixing by H1N1 IAV exhibited a sigmoidal dependence on receptor density when plotted on a log_2_ scale for both receptor types. Lipid-mixing dependence on receptor saturation differed among the mutants. The high-avidity E246G mutant exhibited enhanced lipid mixing at the lowest receptor densities for both receptor types. The double mutant E246G/A227T showed an intermediate phenotype, approaching E246G on 3’SL liposomes and A227T on 6’SL liposomes. Surprisingly, A227T most closely resembled the WT parent virus in receptor dependence for both receptor types. There were no clear effects on lipid-mixing lag time from HA1 mutation, receptor density, or receptor type, though these measurements were highly variable (Fig. S13*B*, *left*). Together, these results support HA-receptor interactions as determinants of lipid-mixing efficiency, with increased receptor avidity promoting fusion within specific receptor-density regimes. They further reveal substantial complexity in how HA point mutations alter lipid-mixing outcomes across receptor types and density.

### Receptor binding promotes HA insertion

We considered two mechanistic explanations for HA-receptor interactions promoting lipid mixing. HA-receptor binding might drive forward the low pH-triggered conformational changes by either prolonging HA1 dilation or promoting HA2 extension (Fig. 1*A*). Alternatively, HA-receptor binding might increase the probability of HA insertion over nonproductive refolding (Fig. 1*C*, *left*) without affecting the preceding pH-dependent transitions. In the first scenario, increased receptor density or NAI would shift the pH dependence of lipid-mixing efficiency toward higher pH values without influencing the magnitude of the change with pH. In the latter scenario, increased receptor density or NAI would not alter the pH dependence of lipid mixing but would instead increase its efficiency across pHs.

To assess the underlying mechanism of fusion promotion by HA-receptor interactions, we performed spCALM with H3N2 IAV and 6’SL-receptor liposomes at 5% or 60% saturation in the presence of absence of 100 nM NAI over a pH range from 4.8 to 7.4 (Fig. 5*A*,*B*). We observed the expected inverse sigmoidal relationship between pH and lipid-mixing efficiency for all conditions (Fig. 5*A*). Notably, an F-test indicated that fitting all curves with a shared IC50 was preferable to a model allowing different IC50 values (p = 0.8157), indicating that neither receptor density nor NAI feeds into the pH-dependent pathway. In contrast, lipid-mixing efficiency was markedly reduced for the low receptor condition without NAI (5% receptor, vehicle) compared to conditions with higher receptor densities, consistent with HA-receptor binding increasing the probability of HA insertion over inactivation. We additionally performed gamma cumulative distribution fitting on the lipid-mixing kinetic trajectories and found no difference in the number of underlying rate-limiting steps between the tested receptor conditions, suggesting that neither reduced receptor density nor NA activity fundamentally alters the reaction stoichiometry (Fig. S8). Virion-liposome binding was more efficient for 60% versus 5% receptor density (Fig. S14). While lipid-mixing lag time decreased at lower pHs, neither receptor density nor NAI had a detectable effect on lag time at pHs where measurements were feasible (Fig 5*B*). The differential effects of pH and receptor conditions on lag time reinforce the interpretation that they act through distinct underlying mechanisms.

**Figure 5.**
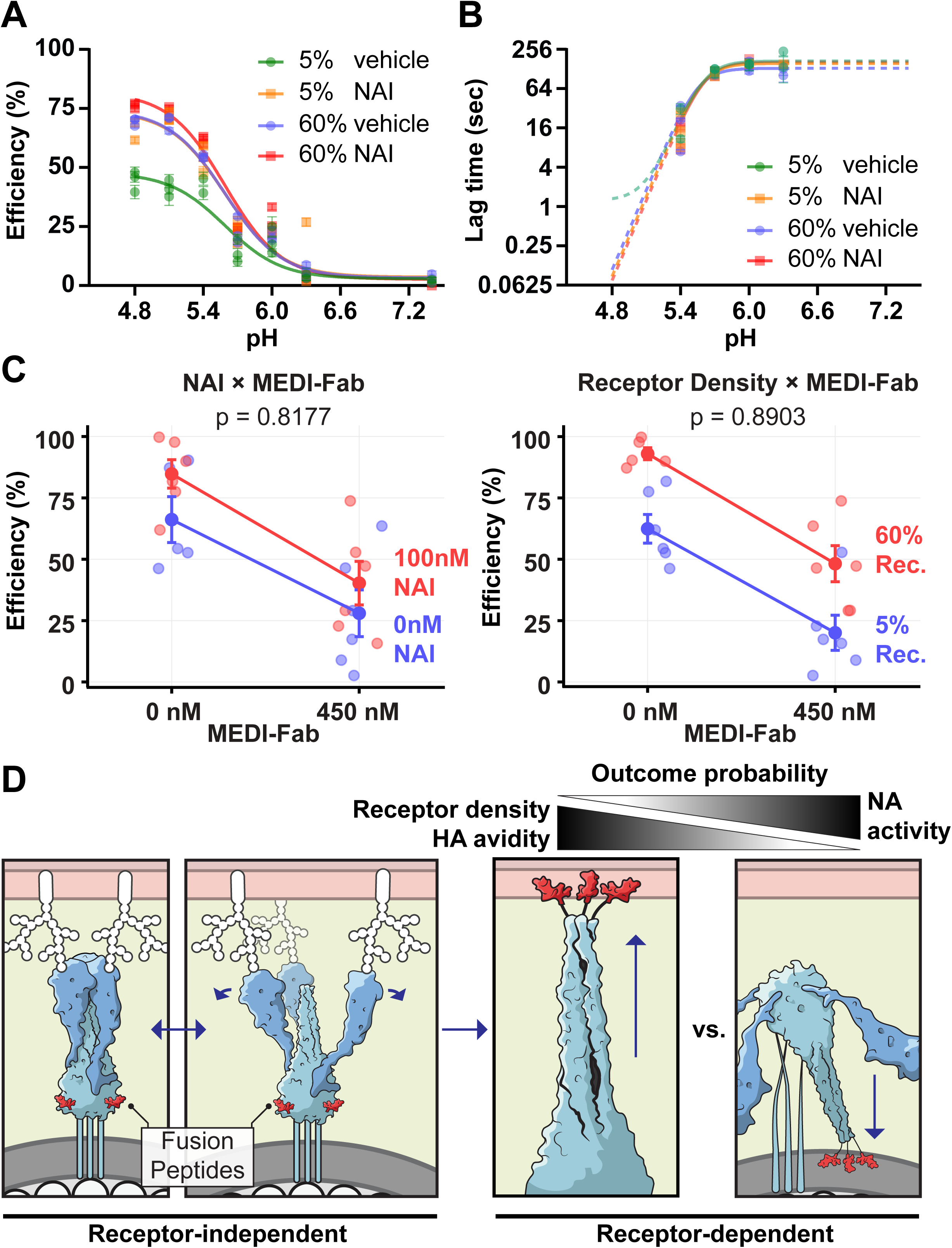
Mechanistic dissection and model: receptor binding promotes fusion-peptide insertion without driving HA extension. (*A*-*B*) Lipid mixing between H3N2 virions and 6’SL-receptor liposomes at 5% or 60% saturation in the presence (NAI) or absence (vehicle) of 100 nM NAI over a range of pHs. The 60% 6’SL receptor saturation (no NAI) condition corresponds to the dataset in Fig. 3E. (*A*) Efficiency. (*B*) Lag time. Plotted are data points from three independent experiments with respective measurement errors. Logistic curves with a shared Hill slope were fit to the combined data. Lag times could not be measured below pH 5.4 because they occurred too quickly relative to the measurement lag or at pH 7.4 because lipid-mixing efficiency was too low; the logistic curves are extrapolated to these domains as dashed lines. (*C*) MEDI’s effect on lipid-mixing efficiency is independent of receptor context. Lipid mixing of H3N2 virions with liposomes displaying 6’SL receptors at 5% or 60% saturation, in the presence or absence of 100 nM NAI and/or 450 nM MEDI. Fusion was triggered by dilution into pH 5.68 buffer. Data points derive from three independent experiments. Experiment-wide interaction effects are shown for NAI and MEDI *(left)* and receptor density and MEDI *(right)*. Error bars indicate the standard error of the mean. P-values for interaction effects were derived from three-way ANOVA (*ns*, not significant) (see Supplementary Table 2). (*D*) Model for the role of sialic acid receptor in IAV membrane fusion. The combined data support a model in which HA-receptor interactions increase the probability of fusion-peptide insertion without affecting the probability or the rate of HA extension.

To further test whether receptor binding modulates HA extension, we asked whether MEDI8852-Fab (MEDI), a stem-binding antibody fragment that inhibits fusion-peptide release (Fig. 1*C*), disproportionately inhibits lipid mixing when receptors are limited (28) (25). We evaluated the effect of a sub-neutralizing dose of MEDI (450 nM) in spCALM with H3N2 IAV and 6’SL-receptor liposomes at 5% or 60% saturation in the presence of absence of 100 nM NAI. MEDI decreased lipid-mixing efficiency, increased lag time, and increased binding across all tested conditions (Figs. 5*C*, S15*A*). Three-way ANOVA revealed no evidence that the effect of MEDI on lipid-mixing efficiency or lag time was dependent on receptor density or NA inhibition (efficiency: p = 0.8903 and p = 0.8177; lag time: p = 0.9647 and p = 0.8816) (Figs 5*C* and S15*B*, Supplementary Table 2). Because MEDI’s effects on lipid mixing were independent of the receptor environment, we conclude that its inhibitory action is mechanistically independent from receptor-mediated promotion of lipid mixing. Together, these data support a model in which HA-receptor binding does not drive extension but instead enhances insertion probability (Fig 5*D*). Accordingly, NA-mediated receptor cleavage favors nonproductive HA foldback, opposing the effects of HA-receptor engagement (Fig 5*D*).

## Discussion

The role of sialic acid receptors in membrane fusion by IAV has long been debated, with proposed functions ranging from passive adhesion to active promotion, inhibition, or even fundamental restructuring of the membrane fusion pathway (30, 34, 35, 37). Here, we demonstrate that the IAV receptor, sialic acid, plays a role beyond attachment by regulating molecular events that govern productive membrane fusion. Specifically, factors such as receptor density, the relative strength of individual HA-receptor interactions, and receptor-cleaving viral NA activity are determinants of fusion efficiency. Given that HA-receptor affinity is a determinant of host specificity and that HA and NA functions are evolutionarily constrained, our study adds a critical dimension to our understanding of the mechanistic constraints on viral evolvability, with implications for pandemic adaptation. These insights were enabled by two key methodological advances: precise control of receptor chemistry and density without perturbing other membrane components, and high-statistical-power quantification of rare virion binding and fusion events by spCALM.

Our study builds upon the existing fusion model (Fig. 1) by defining the previously unresolved role of receptor interactions (Fig. 5D). We show that receptor density and HA-receptor affinity determine lipid-mixing efficiency and support a model in which HA-receptor interactions increase the probability of fusion-peptide insertion. Variation of programmed receptor conditions produced little to no measurable change in the rate of lipid mixing, in agreement with previous single-virion reports (Figs S9, S10, S12, S13, S15, and Supplementary Table 2) (33, 39). Despite measurement uncertainty, this absence of a rate effect was unique to receptor variation, as pH and MEDI-Fab produced the expected increases in lag time for comparable changes in efficiency (Figs. 3*E*, S15, and Supplementary Table 2) (25). Importantly, because lipid-mixing kinetics are largely inconsequential for infectivity whereas reduced fusion efficiency strongly attenuates infection (25), the efficiency changes revealed by spCALM suggest that receptor-mediated effects on fusion might have significant biological consequences.

The mechanism by which receptors favor fusion-peptide insertion remains unclear. One possibility is that HA-receptor interactions stabilize HA in an orientation that favors insertion into the target membrane. HA on virions is predicted to exhibit substantial flexibility, with tilt angles reaching up to ∼50° (46), and receptor binding might constrain this motion. Consistent with this possibility, recent cryoelectron tomography structures of virions bound to sialic-acid receptor mimics revealed receptor-induced HA groupings stabilized in an outward-facing configuration (47). Alternatively, receptor binding might stabilize HA1-HA2 contacts that are present in both the prefusion and extended HA structures but are lost during foldback (19); by favoring extension and delaying foldback, this would increase the probability of insertion over irreversible inactivation. In addition, HA1 and HA2 are disulfide-linked near the trimer base, which rotates by ∼180° during foldback relative to the membrane-inserted domain, pulling HA1 away from the target membrane (48, 49). Stabilization of the target-facing HA1 conformation by receptor binding could therefore impose a free-energy penalty on foldback, facilitating extension and membrane insertion.

Close membrane apposition and subsequent fusion require foldback of 3-4 neighboring HA trimers to overcome the hydration force barrier, estimated to be 30-90 k_B_T (50). Although HA binding to sialyllactose is weak (∼3-7 k_B_T, depending on strain and glycosidic linkage) (18), it is substantial relative to the free energy released from foldback of a single HA trimer (e.g. ∼34.2 k_B_T for HA from X31 H3N2 IAV)(51). Because an HA trimer can engage up to three receptors, foldback free energy could be strongly penalized at high receptor density, potentially increasing the number of HAs required to form a fusion cluster or hindering foldback. However, the unchanged median lag time and shape of lipid-mixing lag-time distributions (N_gamma_ ∼ 3-4 across receptor conditions) (Figs. 5B and S8) across receptor densities tested here (≤ 1 mol%) suggest that receptor binding did not alter fusion-cluster stoichiometry. We thus revealed a regime where receptor binding is a positive regulator of fusion efficiency, where fusion-peptide insertion is enhanced without changes in stoichiometry.

Studies examining fusion in other receptor regimes have reported receptor-induced reconfiguration of the fusion pathway (35) or reduced lipid-mixing kinetics and efficiency (30, 52); however, both findings could be reconciled within the framework we present here. In spCALM, virions are captured in the prefusion state across receptor densities, whereas the study reporting pathway reconfiguration compared receptor-containing to receptor-free membranes, where capture via fusion-peptide insertion likely randomized the initial virion state. Furthermore, the studies reporting inhibition of lipid mixing used much higher receptor densities (10-15 mol%) (30, 52), which might impose a substantial free-energy penalty for receptor unbinding with consequences on foldback. These effects might outweigh enhanced fusion-peptide insertion, dominating reaction kinetics and/or efficiency. Together, these considerations suggest an optimal intermediate receptor density: fusion is limited at low density by inefficient insertion and at high density by foldback hindrance. Future experiments will compare the two extremes of receptor density.

Our results further indicate that NA contributes to the outcome of membrane fusion by cleaving receptors and thereby modifying the receptor environment within the contact patch. It remains unclear whether NA exerts this effect prior to acidification or whether it remains active within the contact patch during membrane fusion. The former scenario is consistent with the clustered NA distribution in regions separate from HA on virions (5, 46, 53), its localized receptor depletion on receptor-bearing surfaces (4, 5, 54), and, for H3N2, its neutral pH-optimum (55). However, several observations favor the latter scenario. Dissociation of NA from the underlying matrix protein M1 at low pH (56) might relieve constraints on NA spatial organization (5), allowing it to diffuse into the contact patch. Furthermore, receptors from the surrounding membrane would likely re-equilibrate with those within the interface. Finally, although NA activity is reduced at low pH for some strains, others have lower pH optima (55), and even suboptimal NA activity might suffice to modulate local HA-receptor interactions on path to fusion.

Independent of the mechanism of NA-mediated fusion antagonism uncovered here, NA inhibitors such as oseltamivir might facilitate fusion in natural infections. Whether NAI inhibits or promotes infection likely also depends on the receptor regime present during infection. Under receptor-rich conditions, or in IAV strains adapted for receptor binding, reduced receptor cleavage might cause impaired progeny virion release to dominate over any fusion benefit, resulting in net inhibition of infection; at high receptor densities, NAI might also inhibit fusion (see preceding paragraph). Consistent with this balance, increased receptor avidity in HA mutants promoted fusion at low receptor densities in spCALM but inversely correlated with plaque size in MDCK cells (Figs. 4*C* and S13*A*). In contrast, under receptor-poor conditions, such as during spillover infection before HA-receptor affinity adapts, NA inhibition might be less effective at impairing virion release but might enhance fusion, thereby promoting infection. If confirmed, this would suggest that antiviral drugs inhibiting NA could inadvertently facilitate viral infection during spillover events. Future experiments measuring the effect of oseltamivir on endosomal membrane fusion in different host cells will test this possibility. In sum, the functional balance between HA and NA determines membrane fusion efficiency, which likely also constrains their evolution.

Multivalent engagement of low-affinity sialic acid receptors by HA is well established to generate high avidity and attachment specificity (18). Our experiments reveal that receptors also play a role after attachment and influence membrane fusion in a manner distinct from their effect on virion attachment. Neutralizing antibodies (Fig. 2C, S15, and Supplementary Table 2) or receptor perturbations (Figs. 4, 5, S9, S10, S12-15, and Supplementary Table 2) produced quantitatively and qualitatively different effects on attachment and membrane fusion, reflecting different underlying dependences. These observations are consistent with our model in which HA-receptor interactions modulate the probability of fusion-peptide insertion by individual HAs within the contact patch, whereas attachment arises from the collective engagement of many HAs with many receptors. Because many viruses rely on weak-affinity glycan receptors for cell entry, the conceptual and experimental framework developed here might find broad applicability.

## Materials and Methods

### Viruses and reagents

Viruses A/PR/8/34 (PR8; H1N1) and A/Udorn/1972 (Udorn) containing X31 HA (XUdorn; H3N2) were generated using an 8-plasmid pHW-based reverse genetics system (57). All WT PR8 pHW plasmids and the X31 HA pHW plasmid were generously provided by Drs. Erich Hoffmann, Richard Webby, and Robert Webster (St. Jude). Site-directed mutagenesis was performed on the PR8 HA pHW plasmid to generate the HA1 binding avidity mutations E246G, A227T, and E246G/A227T. For XUdorn, the remaining seven Udorn gene segments were synthesized and cloned into the pHW backbone. Infectivity in plaque-forming units (PFU)/ml was measured by plaque assay as described previously (20). For virus rescue, the eight pHW plasmids corresponding to each viral genome segment were co-transfected into co-cultures of MDCK-Siat1 and 293T cells. Rescued PR8 viruses were passaged twice at low multiplicity (MOI = 0.002 PFU/cell) then amplified at high multiplicity (MOI = 12 PFU/cell) in MDCK-Siat1 cells (Sigma). Rescued XUdorn virus was passaged twice at low multiplicity (MOI = 0.002 PFU/cell) then amplified at high multiplicity (MOI = 3 PFU/cell) in Calu3 cells.

Viruses were purified and fractionated as described previously (58). Briefly, virus from the final infection supernatant was concentrated by ultracentrifugation at 100,000 x g for 1 hour, purified through a 20% (w/v) sucrose cushion, then the monodisperse, spherical virus fraction was collected on a 20-60% continuous sucrose gradient. Recent protocol modifications designed to minimize particle loss and aggregation were implemented in the case of XUdorn virus preparation as follows. The buffers omitted EDTA, all tubes were pretreated with 0.1% bovine serum albumin (BSA) and rinsed with PBS, and all virus pelleting steps included a 60% layer to avoid pelleting the virus onto the tube surface directly. Expected particle shape was confirmed and counts were determined by flow virometry as reported in (59), and total protein concentration was measured by BCA assay (Thermo Scientific).

For a full list of reagents with source information, tissue culture methods, and antibody preparation information, see **SI Methods**.

### Synthesis of receptor-ssDNA

To synthesize receptor-ssDNA, a two-step approach was used (Fig. 3*C*). First, azido-sialyllactose (**1**) was prepared by reacting commercially available sialyllactose with 2-chloro-1,3-dimethylimidazolinium (DMC) and sodium azide (N_3_) in water, following the method reported by Niu et al (43). Subsequently, **1** was conjugated to 5’-hexynyl-modified ssDNA, which are single-stranded DNAs bearing a terminal alkyne at the 5’-end, to afford receptor-ssDNA. For this, copper-catalyzed Huisgen azide–alkyne cycloaddition (CuAAC) was carried out using copper (I) bromide as the copper source and tris(benzyltriazolylmethyl)amine (TBTA) as a stabilizing ligand. This reaction resulted in an approximate yield of 80%, as determined by denaturing urea polyacrylamide gel electrophoresis (PAGE) analysis (Fig. S5*A-C*). Successful formation of receptor-ssDNA was further confirmed by matrix assisted laser desorption/ionization time of flight (MALDI-TOF MS) (Fig. S5*D,E*). Quantification of the purified conjugate was performed by UV-Vis spectroscopy at 260 nm using NanoDrop analysis. See **SI methods** for a detailed procedure.

### Synthesis of lipid-ssDNA

Precursor lipid, 1,2-*O*-dioctadecyl-*rac*-glycerol, was introduced to 2-cyanoethyl *N,N*-diisopropylchlorophosphoramidite to yield a lipid phosphoramidite **2** (Fig. 3*D*). Specifically, 0.5 g of 1,2-O-Dioctadecyl-rac-glycerol (0.84 mmol) and 320 µL of DIPEA (1.8 mmol) were mixed with 10 mL anhydrous DCM at 0 °C under nitrogen. Then, 0.25 g 2-cyanoethyl *N,N* - diisopropylchlorophosphoramidite (1.3 mmol) was added dropwise to the reaction, which was stirred in an ice bath for 15 minutes before it was returned to room temperature and stirred for another 2 hours. After thin-layer chromatography analysis indicated full consumption of the starting material and formation of the product (Hex:EA:Et_3_N = 90:9:1, Rf = 0.37), the reaction was then washed with NaHCO_3_ 0.5 M, 100 mL), the aqueous layer was further extracted with CH_2_Cl_2_ (2x 20 mL). The organic layer was collected, dried over Na_2_SO_4_, and purified by flash column chromatography (Hex:EA:Et_3_N = 90:9:1) to yield 0.52 g (0.65 mmol, 77% yield) of lipid phosphoramidite product. The synthesized lipid phosphoramidite could then be used as the last “base” to be coupled onto the 5’ end of the ssDNA by a DNA synthesizer, before isolation by preparative HPLC. See **SI methods** for a detailed procedure.

### Single-pair Cytometric Analysis of Lipid-Mixing (spCALM)

R18-labeled virions were preincubated with NAI (vehicle Milli-Q water) and/or relevant antibody (vehicle PBS: pH 7.4, 2.4 g/L KH_2_PO_4_, 14.4 g/L Na_2_HPO_4_, 2 g/L KCl, 80 g/L NaCl) at room temperature for one hour before combining with liposomes or erythrocyte ghosts. Liposomes with the embedded lipid-ssDNA were hybridized to premixed receptor-conjugated ssDNA and/or receptor-free ssDNA at room temperature for one hour. The hybridization reactions included NAI and/or antibody whenever relevant. Erythrocyte ghosts were similarly pretreated to avoid dilution of treatment in the reactions. See **SI Methods** for procedures for R18 labeling and liposome/ghost preparation.

Unless otherwise indicated, spCALM sampling was performed as follows: virions were added to 1e6 liposomes or ghosts at a virion-to-target ratio of 0.15 in a total volume of 16 μL and incubated at 34 °C with 1100 rpm shaking for five minutes (Thermomixer 5350; Eppendorf). HN20C (20 mM HEPES pH 7.4, 140 mM NaCl, 10 μM CaCl_2_), with or without NAI or antibody treatment, was then added to increase the volume to 124 μL to allow for sampling, and the sample was loaded into the cytometer sample tube loader preheated to 34 °C (Cytoflex S; Beckman Coulter). Tube loader temperature was maintained using a wrap-around heating strip adjusted to 34 °C (TLK-H heater and TC300B controller; Thorlabs). spCALM data acquisition was initiated and sample flow rate held constant at 150 μL/min. After 10 seconds, a 10-fold excess volume of pH-adjusted MAcC buffer (30mM MES, 30mM acetic acid, 140mM NaCl, 10μM CaCl_2_) with or without NAI and/or antibody was added to the sample tube and mixed. Acquisition continued for a total of six minutes. The sample probe was rinsed between samples by flowing 30μL of HN20C buffer at 150 μL/min.

### Data analysis

FlowJo software (v10.9.0; Becton, Dickinson & Company) was used to gate and export single virion-target pair measurements (Figs. 2A, S1, S7) and to quantify virion-target attachment (e.g. Fig 2*D*, *left*). Lipid-mixing efficiency (e.g. Fig 2*D*, *right)* and kinetics (e.g. Fig. S3) were derived from gated data using custom Matlab (R2023a; The Mathworks) software. A detailed description of this analysis procedure can be found in **SI Methods**. Dose-response fitting (linear, four-parameter logistic, and gamma cumulative distribution function) and further statistical analyses (ANOVA, multiple comparisons testing) were performed using Graphpad Prism software (v10.6.1; GraphPad Software). Figures were generated using Matlab, RStudio (2025.05.1+513; Posit PBC), and Graphpad Prism, and arranged in Adobe Illustrator.

## Acknowledgments

We thank Austin J. Athman and Alex Stewart of the Research Technologies Branch (RTB), NIAID, for their illustrations shown in figures 1 and 5D. We thank Harry Fruchtman for his work cloning plasmids coding for HA binding avidity mutants. We thank Edward A. Partlow III for working to generate humanized H36-26 IgG1.

This research was supported in part by the Intramural Research Program of the National Institutes of Health (NIH). The contributions of the NIH author(s) are considered Works of the United States Government. The findings and conclusions presented in this paper are those of the author(s) and do not necessarily reflect the views of the NIH or the U.S. Department of Health and Human Services.

This publication is based on research supported by The G. Harold and Leila Y. Mathers Charitable Foundation (T.I.). We acknowledge support from an NIH Director’s New Innovator Award: 1DP2HG011027-01 (J.N.). The funders had no role in study design, data collection and analysis, decision to publish, or preparation of the manuscript.

---

**Movie 1: Real-time distributional dynamics of R18-labeled H3N2 virions bound to erythrocyte ghosts at pH 5.54 and 34°C.** Pseudocolor scatter (left) and contour (right) plots display R18 fluorescence (y-axis, viral membrane dye) versus blue-laser side scatter (x-axis, particle size) over time. 10 minutes of sample acquisition are represented at 200x playback speed. Each additional contour encloses another 2% of the data in the frame.

**Movie 2: Real-time distributional dynamics of R18-labeled H3N2 virions bound to receptor-displaying liposomes at pH 5.54 and 34°C.** Pseudocolor scatter (left) and contour (right) plots display R18 fluorescence (y-axis, viral membrane dye) versus violet-laser side scatter (x-axis, particle size) over time. 10 minutes of sample acquisition are represented at 200x playback speed. Each additional contour encloses another 2% of the data in the frame.

## Notes

### Competing Interest Statement

The authors have declared no competing interest.

