## Supplementary Appendix for "Influenza A virus membrane fusion is regulated by the balance between receptor binding and cleavage"

**This PDF file includes:**

Supplementary Materials and Methods
Supplementary Figures S1 to S15
Supplementary Tables 1 and 2
Dataset 1: Sequencing of H36-26 Hybridoma Variable Region
Dataset 2: H36-26 (human IgG1) Sequence
SI References

**Other supporting materials for this manuscript include the following:**

Movies 1 and 2
MATLAB code for spCALM analysis (https://github.com/planitzersd/spCALM-analysis.git)

### Supplementary Materials and Methods

**Cells:**

Calu3 cells (ATCC) were propagated in EMEM supplemented with 20% FBS. Madin-Darby canine kidney (MDCK)-2,6-sialtransferase (SIAT1) cells (Sigma-Aldrich) and adherent human embryonic kidney (HEK293T) cells were propagated in DMEM supplemented with 10% FBS. Suspension human embryonic kidney 293F (HEK293F) cells were a gift from S. C. Harrison, Harvard Medical School. HEK293F cells were maintained in FreeStyle 293 Expression Medium (Thermo Fisher Scientific). All infections were performed in OptiMEM (Thermo Fisher Scientific) with 1 μg $ml^{-1}$ TPCK-trypsin (Sigma-Aldrich).

**Antibodies:**

The expression vectors (modified pVRC8400) for MEDI8852 Fab (M-Fab) light and heavy chains were a gift from S. C. Harrison, Harvard Medical School. M-Fab was produced by transient transfection of HEK293F cells with polyethylenimine (1 μg DNA, 1.5 μg PEI, 1E6 cells) as described previously (1). A hybridoma producing anti-HA head antibody H36-26 (HA1 Sb epitope) was a gift from J. W. Yewdell (NIH/NIAID/LVD/CBS). Hybridoma cells were sent to BioIntron for variable region sequencing (SI Dataset 1) and synthesis of a plasmid expressing the humanized H36-26 IgG1 (SI Dataset 2).

**ssDNA reagent synthesis and characterization procedure:**

All reactions were performed under ambient atmosphere unless otherwise specified.

*Azide-Functionalized Sialyllactose (1) Synthesis*

2-chloro-1,3-dimethylimidazolinium chloride (16 mg, 0.095 mmol, 3 equiv) was added to a mixture of 6'-sialyllactose (20 mg, 0.031 mmol) (Biosynth), *N,N*-diisopropylethylamine (56 μL, 0.315 mmol), and sodium azide (20 mg, 0.315 mmol) in deuterium oxide (220 μL) ($D_2O$), maintained in an ice bath. (Fig. 3*C*). The reaction mixture was stirred in the ice bath for 1 h. The mixture was then concentrated under reduced pressure and the residue was dissolved in *N,N*-dimethylformamide. The solid was removed by filtration and the filtrate was again concentrated. The residue was dissolved in water and washed with dichloromethane. Purification by ion-exchange column chromatography (Amberlite IR-120B, $Na^+$ form), followed by lyophilization yielded 6′-SL–$N_3$ (18.2 mg, 0.028 mmol, 88%) as a white solid. The NMR data was in good agreement with those published in the literature by Niu et al.[1] $^1$H NMR (500 MHz, Deuterium Oxide) δ 4.79 (d, J = 8.9, 1.2 Hz, 1H), 4.44 (d, J = 7.9 Hz, 1H), 4.02 – 3.51 (m, 18H), 3.39 – 3.34 (t, 1H), 2.77 – 2.68 (dd, 1H), 2.07 – 2.00 (s, 3H), 1.74 (t, J = 12.2 Hz, 1H). HRMS (ESI): Calcd. for $C_{23}H_{37}N_4O_{18}$ $[M+Na]^+$: 681.5491; found: 681.5320.

*Copper-catalyzed Huisgen azide–alkyne cycloaddition (CuAAC) conjugation of 1 to ssDNA*

5′-Hexynyl-ssDNA (90 μL, 100 μM stock in water) and **1** (9 μL, 100 mM stock in water) were combined in a 1.5 mL microcentrifuge tube. The tube cap was removed and replaced with a rubber septum, and the reaction mixture was purged with nitrogen for 15 min. The CuAAC reaction was initiated by addition of 90 μL of a 2 mM premixed Cu/TBTA catalyst solution (1:1 molar ratio). The catalyst stock was prepared by dissolving CuBr (1 mg) and tris(benzyltriazolylmethyl)amine (TBTA, 5.3 mg) in 3.5 mL of a 4:3:1 (v/v/v) mixture of water/DMSO/tBuOH. After addition of the catalyst solution, the reaction mixture was purged with nitrogen for an additional 5 min and then incubated for 12 h. Following completion of the reaction, sodium acetate (19 μL, 3 M, pH 5.2) and ethanol (567 μL, 100%) were added to precipitate the DNA conjugate. The mixture was frozen at −80 °C for 30 min and centrifuged at 21,000 × g for 30 min at 4 °C. The resulting pellet was washed once with cold 70% (v/v) ethanol in water, then dissolved in water (100 μL) and purified by HPLC.

Reverse-phase HPLC purification was performed on an Agilent 1260 Infinity II system equipped with a Prep-C18 column (Agilent, 250 × 10.0 mm, 5 μm particle size, 100 Å pore size). Elution was carried out using a gradient from 95% 0.1 M triethylammonium acetate (TEAA)/5% acetonitrile to 5% 0.1 M TEAA/95% acetonitrile over 30 min. Under these conditions, typical conversion of 5′-hexynyl-ssDNA to receptor-ssDNA was approximately 80%, as determined by denaturing urea polyacrylamide gel electrophoresis (PAGE). The identity of the receptor-ssDNA was further confirmed by matrix-assisted laser desorption/ionization time-of-flight mass spectrometry (MALDI–TOF MS) (Figure S5).

*Synthesis of Lipid Phosphoramidite (2)*

The preparation of lipid-ssDNA started from the introduction of the lipid, 1,2-*O*-dioctadecyl-*rac*-glycerol, to 2-cyanoethyl *N,N*-diisopropylchlorophosphoramidite to give a lipid phosphoramidite **2** (Fig. 3D). Specifically, 0.5 g of 1,2-O-Dioctadecyl-rac-glycerol (0.84 mmol) and 320 µL of DIPEA (1.8 mmol) were mixed with 10 mL anhydrous DCM at 0 °C under nitrogen. Then, 0.25 g 2-cyanoethyl *N,N* -diisopropylchlorophosphoramidite (1.3 mmol) was added dropwise to the reaction, which was stirred in ice bath for 15 minutes before being returned to room temperature and stirred for another 2 hours. After TLC indicated full consumption of the starting material and formation of product spot (Hex:EA:$Et_3N$ = 90:9:1, Rf = 0.37), the reaction was then washed with $NaHCO_3$ 0.5 M, 1 00 mL) and the aqueous layer was further extracted with $CH_2Cl_2$ (2x 20 mL). The organic layer was collected and dried over $Na_2SO_4$, evaporated to dryness, and purified by flash column chromatography (Hex:EA:$Et_3N$ = 90:9:1) to yield 0.52 g (0.65 mmol, 77% yield) of lipid phosphoramidite product. The synthesized lipid phosphoramidite can then be used as the last "base" to be coupled onto the 5' end of the ssDNA by DNA synthesizer, before isolation by preparative HPLC.

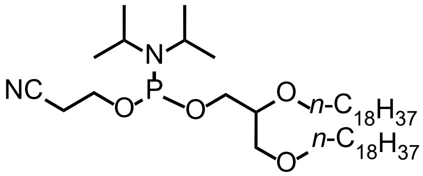


**$^1$H NMR** (600 MHz, $CDCl_3$) δ 3.90 – 3.40 (m, 13H), 2.63 (dq, *J* = 6.3, 3.1 Hz, 2H), 1.58 – 1.52 (m, 4H), 1.35 – 1.22 (s, 60H), 1.21 – 1.14 (m, 12H), 0.88 (t, *J* = 7.0 Hz, 6H). **$^{31}$P NMR** (202 MHz, $CDCl_3$) δ 148.5 (d, J = 30.0 Hz).

*Synthesis of Lipid-ssDNA*

The oligonucleotides used in this study were synthesized with an in-house Mermade6 DNA/RNA synthesizer using standard phosphoramidite chemistry (2). Reagents for solid phase synthesis were purchased from Glen Research including the phosphoramidite monomers, activator, deprotection reagent and capping reagents. The resulting oligonucleotides were subsequently deprotected using AMA (conc. Ammonia / 40% Aqueous Methylamine, V / V = 1:1) and purified using Glen-Pak™ cartridge (Glen Research, 60-5100) following the manufacturer's protocol. The target products were then lyophilized, dissolved with water and characterized by Bruker Autoflex LRF Speed mass spectrometer.

*Characterization of reaction products:*

1H and 13C were collected in $D_2O$, unless otherwise noted, on a 600 MHz Varian/Agilent spectrometer or 500 MHz VNMRS Varian/Agilent spectrometer or 500 MHz Bruker system with a helium CryoProbe and autosampler while using residual water peak (δ = 4.79 for H) as an internal standard.

High-resolution mass spectrometry was performed on a JEOL AccuTOF-DART (positive mode) or Agilent 6220 TOF-ESI (positive mode). Matrix-Assisted Laser Desorption/Ionization Time-of-Flight Mass Spectrometry (MALDI-TOF MS) was performed on a Bruker Auto Flex Max instrument (negative mode) while 3-hydroxypropionic acid (3-HPA) was used as matrix with diammonium citrate added as the counterion source.

High-performance liquid chromatography (HPLC) was carried out with an Agilent HPLC system 1260 Infinity II LC System containing an Agilent 5 Prep-C18 (250 x 10.0 mm 5 µM column with 100 Å packing material) HPLC column at 25°C and a flow rate of 2 mL/min. The eluent used was triethylammonium acetate (TEAA) (0.1 M pH 7.0) and HPLC grade acetonitrile. Absorbance values were collected on a Thermo Scientific™ NanoDrop™ One/OneC Microvolume UV-Vis Spectrophotometer.

**Virion R18 labeling:**

Rhodamine B chloride (R18, Invitrogen), dissolved in 200 proof ethanol, was diluted in HN20C buffer and added to a suspension of virions such that the final ethanol concentration was 1% vol/vol and viral protein concentration was 0.24 mg/mL. PR8 WT (H1N1), E246G, A227T, and E246G/A227T were labeled with 10 µM R18, and XUdorn (H3N2) was labeled with 4 µM R18. The R18-virus mixture was incubated at room temperature, in the dark, for one hour. To exclude aggregates, H1N1 labeling reactions were centrifuged at 1000 x g for 2 hours in BSA-treated low-binding microcentrifuge tubes and the supernatants were carefully harvested to exclude aggregates; this step was omitted for H3N2. The labeled viruses were split into single-use aliquots in PCR tubes and stored frozen at -80 °C. For an experiment, aliquots were thawed quickly and used the same day. Virus particle concentrations were obtained by flow virometry after thawing as described in (3).

**Preparation of human erythrocyte ghosts:**

A published protocol for generating bovine erythrocyte ghosts (4) was adapted for human red blood cells used here: irradiated, leukocyte-reduced human red blood cells treated with ACD-A and AS-1 were procured from the NIH Blood Bank (Bethesda MD, USA). Cells were washed in cold PBS with 2 mM magnesium chloride (PBS-Mg) at 4300 rcf until the supernatant clarified. Cells were then cycled between suspension in PBS-Mg and lysis buffer (5 mM sodium phosphate, pH 7) and spun between steps at 14,900 rcf; wash cycles were continued until the cells formed a fluffy white pellet with a small amount of residual reddish cells at the tube bottom and with clear, colorless supernatant. The pellet was transferred to a new tube and spun at 16490 rcf. The topmost fluffy white layer of the pellet was harvested, excluding any reddish cells at the bottom. The harvested cells were then suspended in two pellet volumes of lysis buffer, to which 1/9 pellet volume of the 10X resealing buffer (1.2 M KCl, 300 mM NaCl, 10 mM MgCl2, 100 mM sodium phosphate, pH 7) was added. The cells were incubated on ice for 15 minutes and then at 37 °C for 60 minutes. The cells were washed twice in PBS-Mg at 16490 rcf, stored at 4°C in a plastic-sealed microcentrifuge tube filled with ultrapure argon gas, and used within 5 months.

**Desialylation of erythrocyte ghosts using bacterial neuraminidase**

20µL of 1.2e7 cells/µL ghosts in PBS-Mg were incubated in 1 ml cleavage buffer (50 mM acetate pH 5.5, 150 mM NaCl, 4 mM $CaCl_2$) with or without 32 µg/mL cholera filtrate at 37 °C with shaking at 400 rpm for 10 minutes; the reactions were then moved to ice. 10 µL of the reaction was reserved for lectin-staining analysis; the rest was washed three times with 1mL of cold PBS-Mg, spun at 16000 rcf for 10 minutes, and then resuspended in 10-20 µL of ice-cold PBS-Mg. *Sambucas nigra* lectin (SNA/EBL, Vector Laboratories) was conjugated in-house to Janelia Fluor® 646 NHS ester (Tocris) at a 20:1 dye:protein ratio. A sample of the neuraminidase reaction was washed once with lectin buffer (10 mM HEPES pH 8.51, 150 mM NaCl, 0.1 mM $CaCl_2$) and then incubated with 5.5 nM JF646-SNA lectin in lectin buffer for 15 minutes at room temperature. The cell suspension was then diluted 10-fold in lectin buffer. JF646-fluorescence signal on mock versus treated ghosts was measured by flow cytometry (Fig. S4).

**Preparation of DNA-functionalized liposomes:**

A solution with a 40 : 40 : 20 : 0.025 : 0.0025 : 0.0452 molar ratio of chloroform-dissolved DOPC (1,2-dioleoyl-sn-glycero-3-phosphocholine, Avanti Research), POPC (1-palmitoyl-2-oleoyl-glycero-3-phosphocholine, Avanti Research), cholesterol (Avanti Research), fluorescein-DHPE (N-(Fluorescein-5-Thiocarbamoyl)-1,2-Dihexadecanoyl-sn-Glycero-3-Phosphoethanolamine, Triethylammonium Salt, Invitrogen), and biotin-PE (1,2-dioleoyl-sn-glycero-3-phosphoethanolamine-N-(cap biotinyl) (sodium salt), Avanti Research) and ethanol-dissolved DiD (1,1'-Dioctadecyl-3,3,3',3'-Tetramethylindodicarbocyanine, 4-Chlorobenzenesulfonate Salt, Invitrogen) was prepared and dried into a thin film under a stream of argon gas in a borosilicate test tube. The film was desiccated under house vacuum for at least two hours, or overnight. A solution of lipid-ssDNA in HN20C buffer was used to resuspend the dried film to a total lipid concentration of 9.24 mg/mL with 1 mol% of lipid-ssDNA by vortexing and shaking at 37 °C for 30 minutes (Infors Multitron incubator). The suspended lipids were then subjected to five cycles of vortex mixing, rapid freezing in liquid nitrogen, and rapid thawing in warm water. The crude suspension was homogenized by extruding (Avanti Research) 30 times through a 1 µm filter

(Whatman). The liposomes were degassed under house vacuum for 10 minutes, stored at 4 °C under argon in plastic-sealed tubes, and used within two months of preparation.

**Single-particle TIRF microscopy on supported planar bilayers**

Liposomes incorporating 1 mol% ssDNA18-lipid were prepared as described in the preceding section, except that DiD was omitted and a 200 nm filter (Whatman) was used for extrusion. Planar bilayers were formed by spontaneous vesicle spreading in home-built PDMS flow cells on coverglasses as described previously (5). The membranes were then labeled sequentially with 30 µg/ml fluorescein-conjugated streptavidin for 10 minutes and ssDNA18-lipid was hybridized with a 0.3 µM mixture of 3'SL receptor-DNA12 and receptor-free-DNA12, with HNE20 buffer (20mM HEPES pH 7.4, 140mM NaCl, 0.2mM EDTA) wash steps in between. R18-labeled H3N2 virions were flowed for one minute and washed with HNE20 buffer before imaging. The flow cell was imaged using an RM21 Advanced microscope with Micro-Mirror TIRF illumination (Mad City Labs) and 488 nm (2.03 µW) and 552 nm (18.0µW) lasers (Coherent). Emission light was spatially separated using a MadView module (Mad City Labs) and detected on separate regions of the Orca Quest camera sensor (C15550-20UP, Hamamatsu). All microscope systems were controlled using Micro-Manager software (https://micro-manager.org/)

**NA-activity inhibition assay**

Inhibition of NA activity of H3N2 virions by NAI was assessed using the NA-Fluor™ Influenza Neuraminidase Assay Kit (4457091, Invitrogen), which measures cleavage of the fluorogenic substrate MUNANA by viral NA. The assay was performed according to the kit instructions (Fig. S11).

**Data analysis**

The FlowJo gate for bound virion-target pairs was established based on a liposome-only or ghost-only control sample such that occupancy in the pairs gate was ≤ 0.3 % of the total liposomes or ghosts. These single-pair observations were analyzed using custom Matlab scripts to extract the efficiency and lag time of lipid mixing. ChatGPT (GPT-5, OpenAI) was used to accelerate development of analysis procedures. R18-fluorescence values were asinh-transformed, and characteristic reference fluorescence distributions of unfused and lipid-mixed pairs were modeled by pooling a subset of high lipid-mixing efficiency samples then gating the time interval before pH-drop and at reaction plateau when lipid-mixing is complete. The data in each time interval were fit with Burr distributions. In cases where conversion to the fully lipid-mixed state did not occur, the pre-pH drop distribution was used to deconvolve the late time state by gaussian mixture model deconvolution to estimate the characteristic distribution of lipid-mixed pairs. This approach is less prone to underestimating lipid-mixing efficiency than simple R18 fluorescence threshold gating.

For each sample, events were time binned to include approximately the same number of events. The fraction of lipid-mixed pairs in each bin was estimated by computing the posterior responsibility of the lipid-mixed distribution relative to the unfused distribution using an expectation-maximization algorithm. Lipid-mixing efficiency was quantified using a model-agnostic span metric, defined as the difference in mean lipid-mixed fraction between the late-time plateau and pre-pH drop baseline (typically <3%). Uncertainty was estimated by binomial approximation. pH-drop time $t_0$ was estimated by finding the maximum rate of change of detected particle concentration resulting from the low-pH buffer dilution (see Fig. 3*E, right*); uncertainty was determined by bootstrap resampling. Lipid-mixing lag time was estimated by fitting lipid-mixed fraction kinetics with a weighted, nondecreasing function using pooled-adjacent-violators regression. Baseline and plateau levels were estimated as weighted medians of the fit within pre-pH drop (baseline) and plateau time windows, defining a dynamic range $A = plateau - baseline$. Lag time was defined as the first time at which the isotonic fit crossed the midpoint, $baseline + 0.5A$, minus the pH-drop time $t_0$, with linear interpolation between adjacent bins. Error was estimated by bootstrap resampling. Estimates were censored when the inferred dynamic range was not distinguishable from baseline noise, assessed using a MAD-based criterion.

Gamma CDF fitting of lipid-mixing versus time trajectories was performed in Graphpad Prism using the following expression:

$$CDF(t) = Bottom + (Top - Bottom)\frac{\gamma(N,\ k(t-t_0))}{\Gamma(N)}$$

Where $\gamma\big(N, k(t-t_0)\big)$ is the lower incomplete gamma function with shape parameter $N$, rate parameter $k$, and function onset time $t_0$. $\Gamma(N)$ is the gamma function for shape parameter N. $Bottom$ and $Top$ parameters permit starting and ending plateau values greater than zero and less than one, respectively. Gamma CDFs were fit with $t_0$ fixed at the detected time of low pH buffer addition determined during data processing.

### Supplementary Figures

#### Supplementary Figure S1

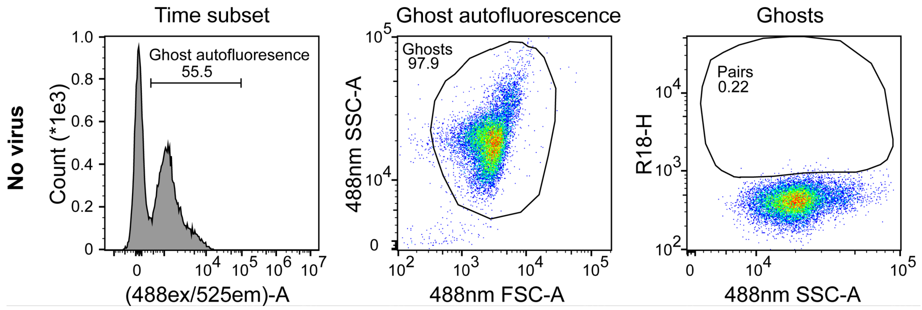


**Fig. S1. The gating strategy for spCALM with erythrocyte ghosts.** Erythrocyte ghosts are identified by autofluorescence in the 488-nm channel (left). Cell singlets are gated on a 488-nm side scatter (y-axis; SSC-A) versus forward scatter (x-axis; FSC-A) plot (middle). Virus-free samples are used to establish the virion-cell pair gate based on viral membrane R18 fluorescence (y-axis; R18-H) and 488nm side-scatter (x-axis; SSC-A), excluding unbound cells (right). Subplot titles indicate the parent gate for the data.

**Supplementary Figure S2**

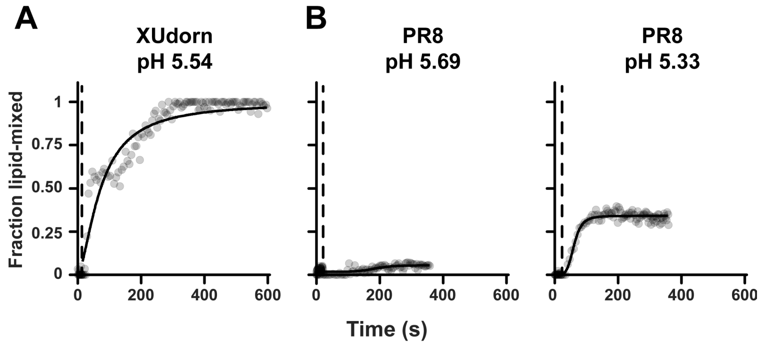


**Fig. S2. Fraction of lipid-mixed pairs as a function of time for H3N2 and H1N1 virions and ghosts.** H3N2 virions exhibit efficient lipid mixing at pH 5.54 (~100% at plateau) at 34 °C, whereas H1N1 requires lower pH. Solid lines are logistic fits; dashed vertical lines indicate the measurement lag.

#### Supplementary Figure S3

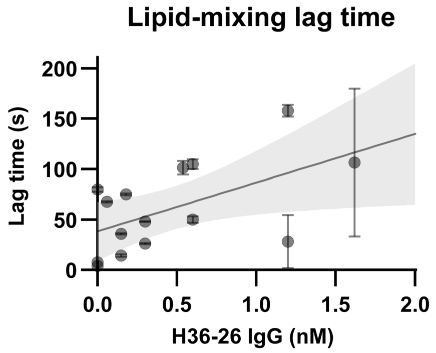


**Fig. S3. Lipid-mixing lag time versus HA1-antibody H36-26 concentration for the experiment in Fig. 2C.** Lag time is defined as the time to the median lipid-mixing extent. Data points derive from three independent experiments; error bars were estimated by bootstrap resampling. The line shows a linear regression with shaded 95% confidence interval.

#### Supplementary Figure S4

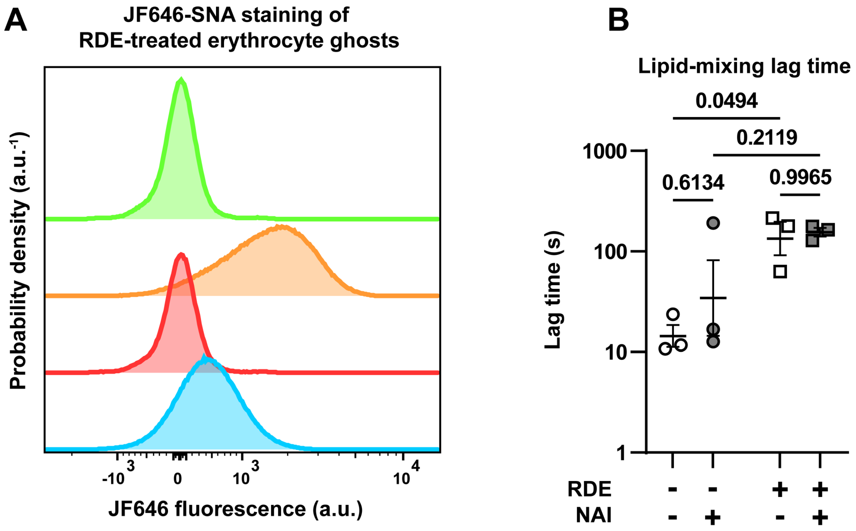


| | Treatment | JF646-SNA | Median fluorescence (a.u.) |
|---|---|---|---|
| (green) | Mock | − | 22.4 |
| (orange) | Mock | + | 1575 |
| (red) | RDE | − | 25.3 |
| (blue) | RDE | + | 454 |

**Fig. S4. Confirmation of erythrocyte desialylation by lectin binding and its effect on lipid-mixing lag time.** (A) JF646 fluorescence distributions for mock- or RDE-treated cells, with or without JF646-SNA lectin. Samples are color-coded and labeled in the table, which also lists the median JF646 fluorescence signal for each sample. RDE treatment reduces JF646-SNA lectin binding to erythrocyte ghosts compared to mock-treatment, consistent with desialylation. (B) Lipid-mixing lag time for the experiment depicted in Fig 2D. Data points deriving from three independent experiments were separately plotted with geometric mean +/- SEM. Statistical significance was determined by Tukey's multiple comparisons test of log-scaled data.

#### Supplementary Figure S5

**A**

| Entry | Copper source (equiv.) | Additives (equiv.) | Solvent (ratio) | $N_3$:Alkyne (ratio) | Result |
|---|---|---|---|---|---|
| 1 | $CuSO_4 \cdot 5H_2O$ (50) | Sodium ascorbate (100) | DMSO:$H_2O$ (56:44) | 100:1 | ≈20% |
| 2 | Cu(I)Br (20) | TBTA (20) | DMSO:$H_2O$:*t*-BuOH (50:46:4) | 100:1 | ≈80% |

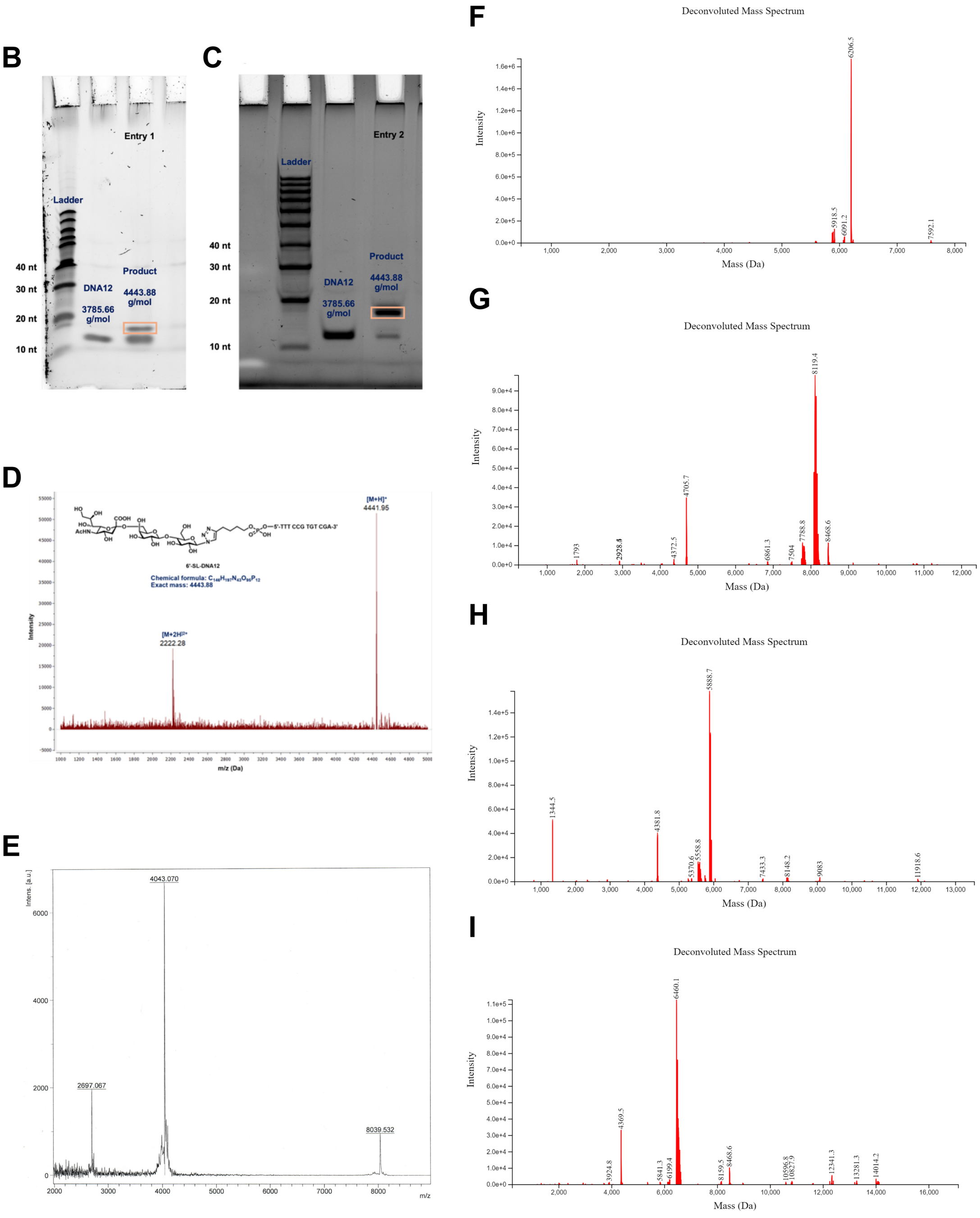


**Fig. S5. Establishment of reaction conditions and characterization of synthesized products.** (A-C) Screening of reaction conditions for the CuAAC conjugation of 6′-SL–$N_3$ with 5′-hexynyl-DNA12. (A) Summary of the reaction conditions evaluated. (B–C) Denaturing urea polyacrylamide gel electrophoresis (PAGE) analysis of the corresponding reactions: (B) entry 1; (C) entry 2. Initial optimization was carried out by evaluating two reaction conditions reported in the literature (2, 3). The parameters examined included the copper source, reductant, stabilizing ligand, and the ratio of cosolvents used to solubilize the reactants (A). The first condition employed copper(II) sulfate pentahydrate with sodium ascorbate as the reductant (A, entry 1). Under these conditions, poor coupling efficiency was observed, with only ~20% conversion as determined by denaturing gel electrophoresis (B). The low conversion is attributed to the use of Cu(II) species, which can interact unfavorably with oligonucleotides and reduce reaction efficiency. To improve the reaction outcome, the copper source was changed to copper(I) bromide in the presence of tris(benzyltriazolylmethyl)amine (TBTA) as a stabilizing ligand (A, entry 2). Under identical conditions and at the same azide-to-alkyne ratio, this modification resulted in a substantial increase in conversion, reaching approximately 80% as determined by gel electrophoresis (C). It was also observed that scaling the reaction beyond 9 nmol of DNA led to decreased conversion efficiency relative to smaller-scale reactions (~1 nmol), for which near-quantitative conversion could be achieved. (D-I) Characterization of synthesized reagents: (D) MALDI–TOF mass spectrum of a receptor-ssDNA 6′SL–DNA12. 3-Hydroxypicolinic acid (3-HPA) was used as the matrix, and diammonium citrate was added as a counterion source. (E) MALDI-TOF mass spectrum of receptor-ssDNA 6’SL-DNA24. The observed m/z of 4043.07 is in close agreement with the expected value of 4044.23. (F) ESI-MS analysis of Lipid-DNA18 (18 nt). The discovered mass (6206.5 Da) agreed with the calculated mass of the desired product (6206.7 Da). (G) ESI-MS analysis of Lipid-DNA24 (24 nt). The discovered mass (8119.4 Da) agreed with the calculated mass of the desired product (8090.9 Da). (H) ESI-MS analysis of Lipid-DNA12XL (17 nt). The discovered mass (5888.7 Da) agreed with the calculated mass of the desired product (5888.5 Da). (I) ESI-MS analysis of Lipid-DNA14XL (19 nt). The discovered mass (6460.1 Da) agreed with the calculated mass of the desired product (6459.8 Da).

#### Supplementary Figure S6

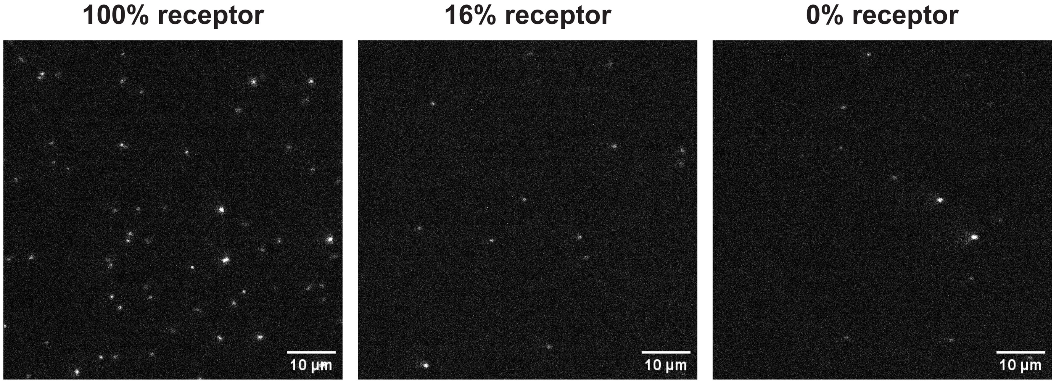


**Fig. S6. Representative TIRF microscopy images of SPBs incorporating 1 mol% lipid-ssDNA18 (see Supplementary Table 1) hybridized to 6'SL-ssDNA12, forming a 12-nt membrane-proximal dsDNA spacer and a 6-nt single-stranded overhang.** R18-labeled H3N2 virions bound to the membrane are excited by the evanescent field of the reflected laser light, appearing as bright, diffraction-limited spots.

#### Supplementary Figure S7

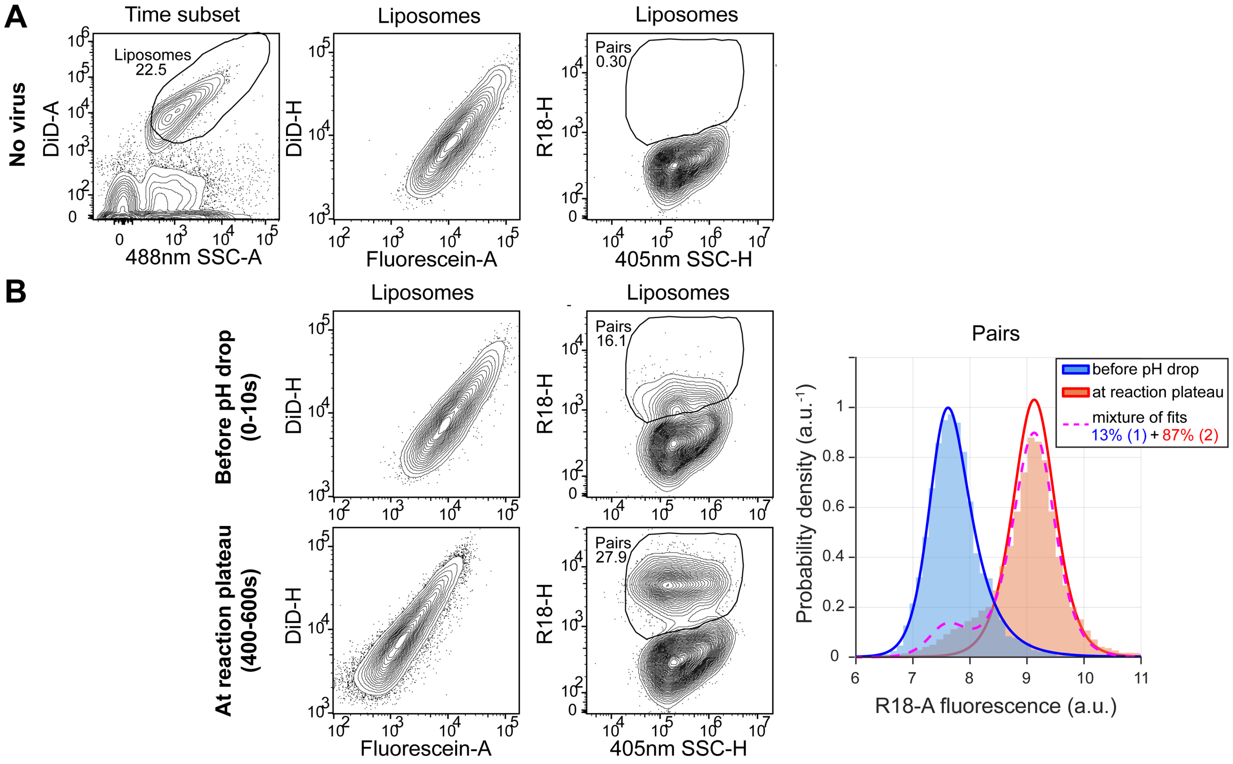


**Fig. S7. The gating strategy for spCALM with liposomes displaying dsDNA-receptors.** (A) Control sample with liposomes and no virus. Liposomes are distinguished from background based on 488nm side scatter (x-axis; SSC-A) and pH-insensitive membrane dye fluorescence (y-axis; DiD-A) (left). pH-sensitive fluorescein fluorescence provides a readout of the reaction pH (middle). No-virus samples help establish the virion-cell pair gate, excluding unbound liposomes (right). Subplot titles indicate the parent gate for the data. (B) Example spCALM reaction windows before pH drop (0-10 s, top) and at the reaction plateau (400-600 s, bottom) for H3N2 virions from Movie 2. pH-drop induces a decrease in pH-sensitive (x-axis; Fluorescein-A) but not pH-stable (y-axis; DiD-H) liposomal dye fluorescence (left). Lipid mixing between virions and liposomes dequenches R18 fluorescence, shifting lipid-mixed pairs to a brighter population distinct from unfused pairs. (middle). Histograms of R18 fluorescence for virion-ghost pairs before acidification (blue) and after the reaction plateau (red), pooled from selected samples which undergo high-efficiency lipid-mixing. Burr distribution fits in each time interval define characteristic unfused and lipid-mixed reference distributions. These distributions are then applied to intermediate time bins to quantify lipid mixing and derive fusion kinetics. Because lipid mixing did not reach completion at the reaction plateau in this experiment, the lipid-mixed distribution was determined by deconvolving the unfused reference distribution from the plateau data.

#### Supplementary Figure S8

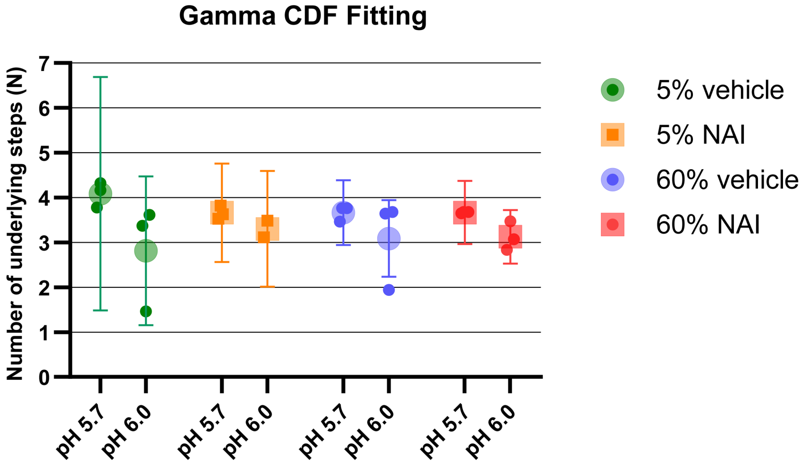


**Fig. S8. N parameter from Gamma cumulative distribution function (CDF) fits to lipid-mixing versus time trajectories at pH 5.7 and 6.0; data from Figs. 3E and 5A**. Data points from three independent experiments are shown separately as solid points; the mean +/- propagated SEM is shown as a larger transparent point with error bars. Measurement lag was used as the onset time for gamma CDF fits.

#### Supplementary Figure S9

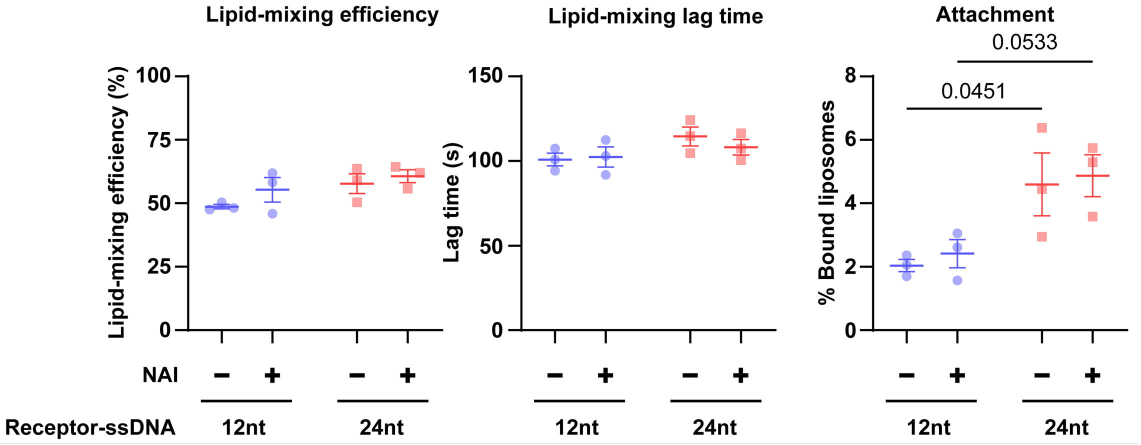


**Fig. S9. Receptor spacer length affects attachment but not lipid-mixing efficiency or lag time.** Lipid-mixing efficiency, lag time, and attachment for H3N2 virions and liposomes displaying 6'SL receptors on 12-nt or 24-nt dsDNA spacers at full saturation. Lipid mixing was triggered by dilution into pH 5.70 buffer in the presence of absence of 100nM NAI. Data points deriving from three independent experiments were plotted with mean +/- SEM. Two-way ANOVA revealed no significant effects of spacer length or NAI-treatment, except for attachment, where spacer length but not NAI was significant (P = 0.0046). Pairwise comparisons between 12-nt and 24-nt spacers within each NAI condition are shown for attachment and were corrected using the Šídák method.

#### Supplementary Figure S10

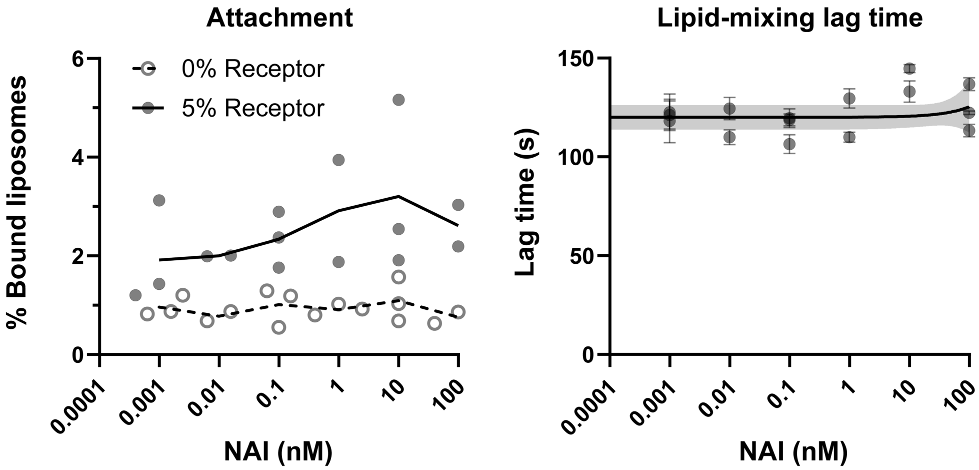


**Fig. S10. Attachment and lipid-mixing lag time for the experiment depicted in Fig 4A.** (Left) Individual data points deriving from three independent experiments were separately plotted with means connected by straight lines. (Right) Individual data points for the 5% receptor condition deriving from three independent experiments were separately plotted; error bars were estimated by bootstrap resampling. A linear regression was fit to the combined data with shaded 95% confidence interval. Lag times could not be determined for the 0% receptor condition due to insufficient lipid mixing.

#### Supplementary Figure S11

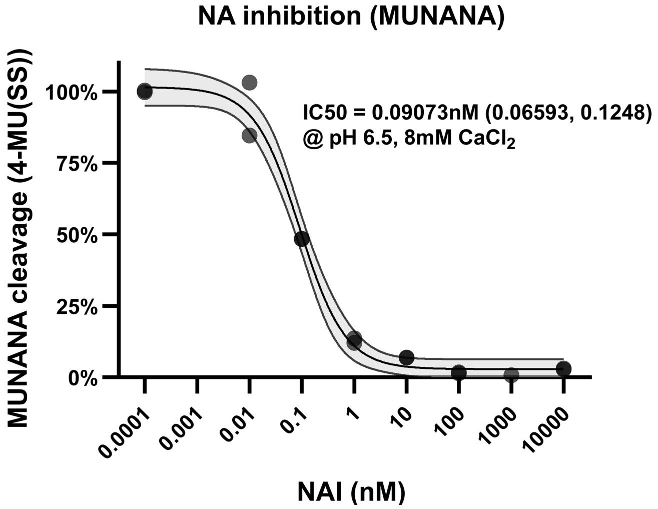


**Fig. S11. NAI dose-response curve for H3N2 NA-activity measured by MUNANA assay.** MUNANA reports virion-associated NA activity via fluorogenic substrate cleavage. Normalized 4-MU(SS) fluorescence is plotted versus NAI concentration. The experiment was performed twice on the same plate. A logistic curve was fit to the combined data with shaded 95% confidence interval. The IC50 and its 95% confidence interval are indicated, along with the assay buffer (33.3mM MES, 8mM CaCl2, pH 6.5).

#### Supplementary Figure S12

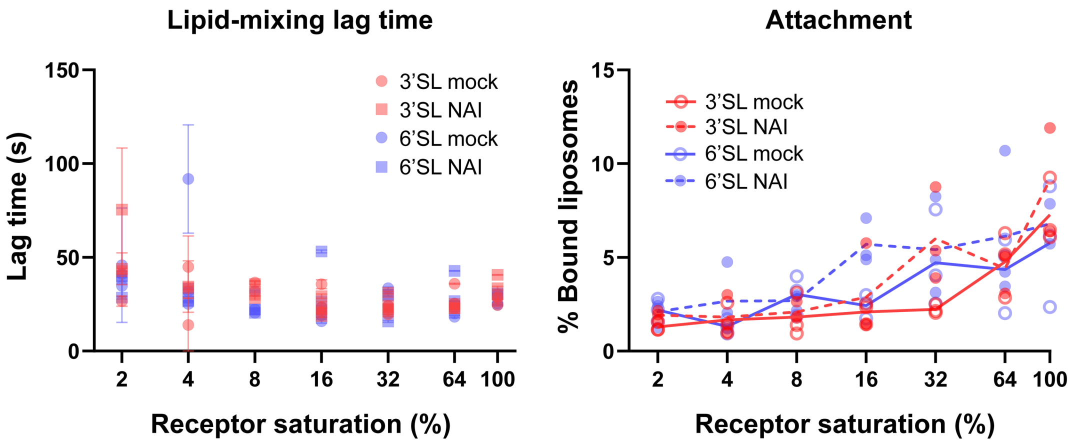


**Fig. S12. Attachment and lipid-mixing lag time for the data in Fig 4B.** (Left) Individual data points deriving from three independent experiments were separately plotted; error was estimated by bootstrap resampling. (Right) Individual data points deriving from three independent experiments were separately plotted with means connected by straight lines.

#### Supplementary Figure S13

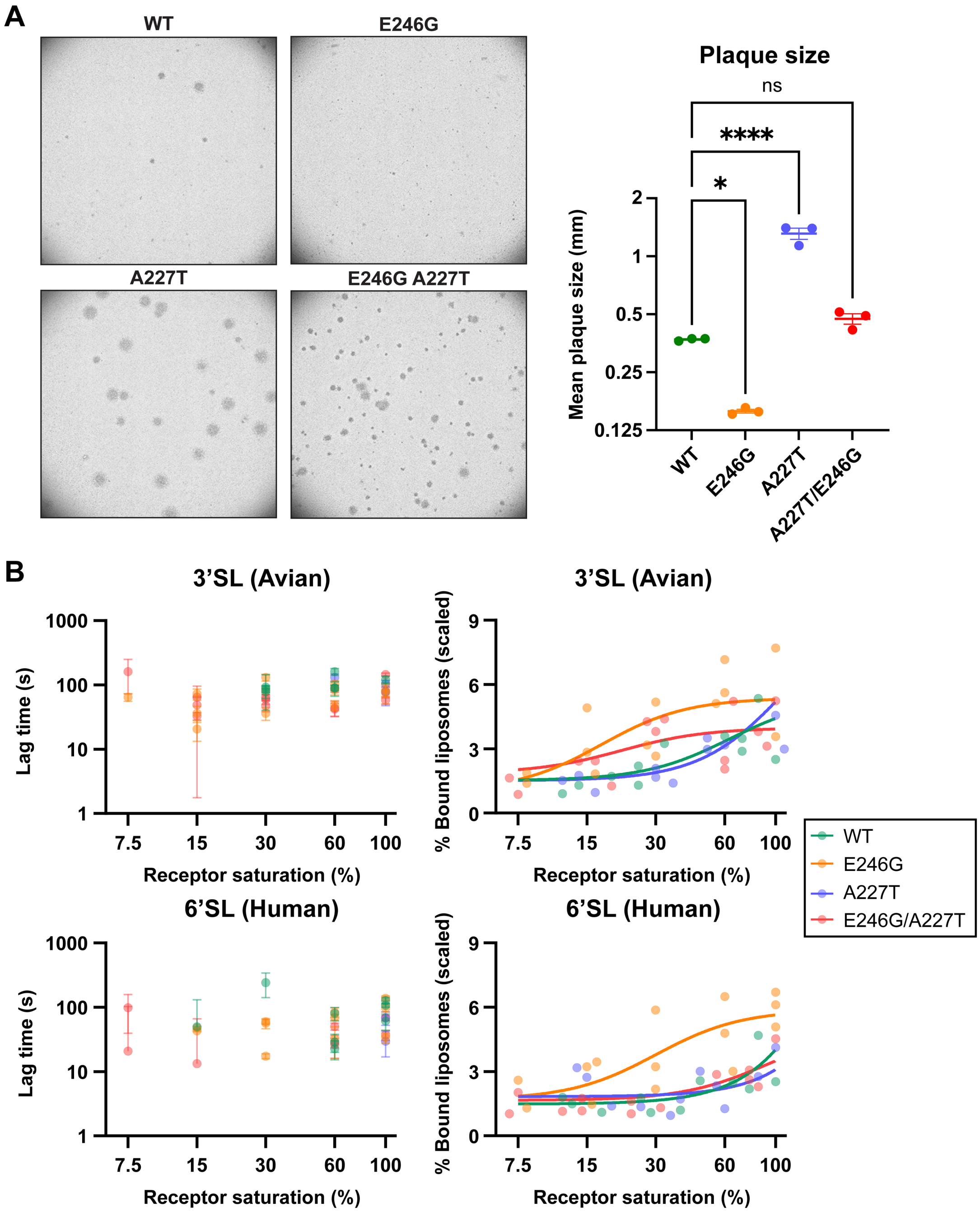


**Fig. S13. Plaque size and spCALM-derived lipid-mixing lag time and attachment of H1N1 WT and HA1 receptor-avidity mutants.** (A) Representative plaque-assay images for H1N1 WT and the E246G, A227T, and E246G/A227T variants (left). Mean plaque sizes from three experiments are shown separately with the grand mean +/- SEM overlaid; n=104, 356, 90, and 341 plaques for WT, E246G, A227T, and E246G/A227T, respectively (right). Statistical significance relative to WT was determined using Dunnet's multiple-comparisons test. ns P >

0.05, *P < 0.05, **P < 0.01, ***P < 0.001, ****P < 0.0001. (B) Lipid-mixing lag time (left) and attachment (right) for the data in Fig. 4C. Top: 3'SL receptor; bottom: 6'SL receptor. Lag-time data deriving from three independent experiments are individually shown except where they could not be reliably determined; error was estimated by bootstrap resampling. Raw attachment values were scaled across virus variants using counts from virus-only input controls. Individual data points from three independent experiments are shown separately. Logistic curves with a shared Hill slope were fit to the combined data.

#### Supplementary Figure S14

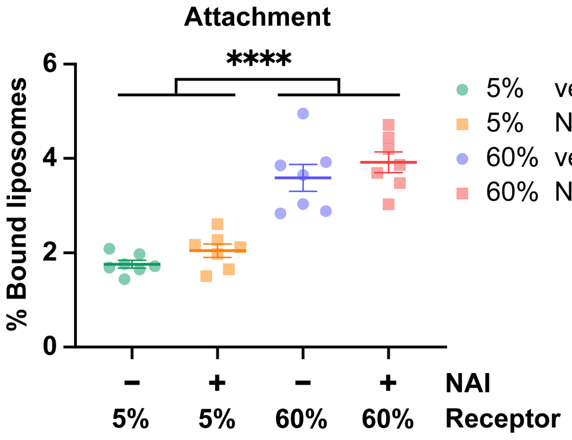


**Fig. S14. Attachment data from the experiment in Figs. 5A-B.** Data from three independent experiments are shown individually with mean +/- SEM. Samples from each receptor/NAI condition were grouped across pH conditions, as attachment was measured before pH drop. Two-way ANOVA indicated that only receptor saturation significantly explained the variance; therefore, only the marginal effect of receptor saturation is shown. ****P < 0.0001 by Tukey's multiple comparisons test.

#### Supplementary Figure S15

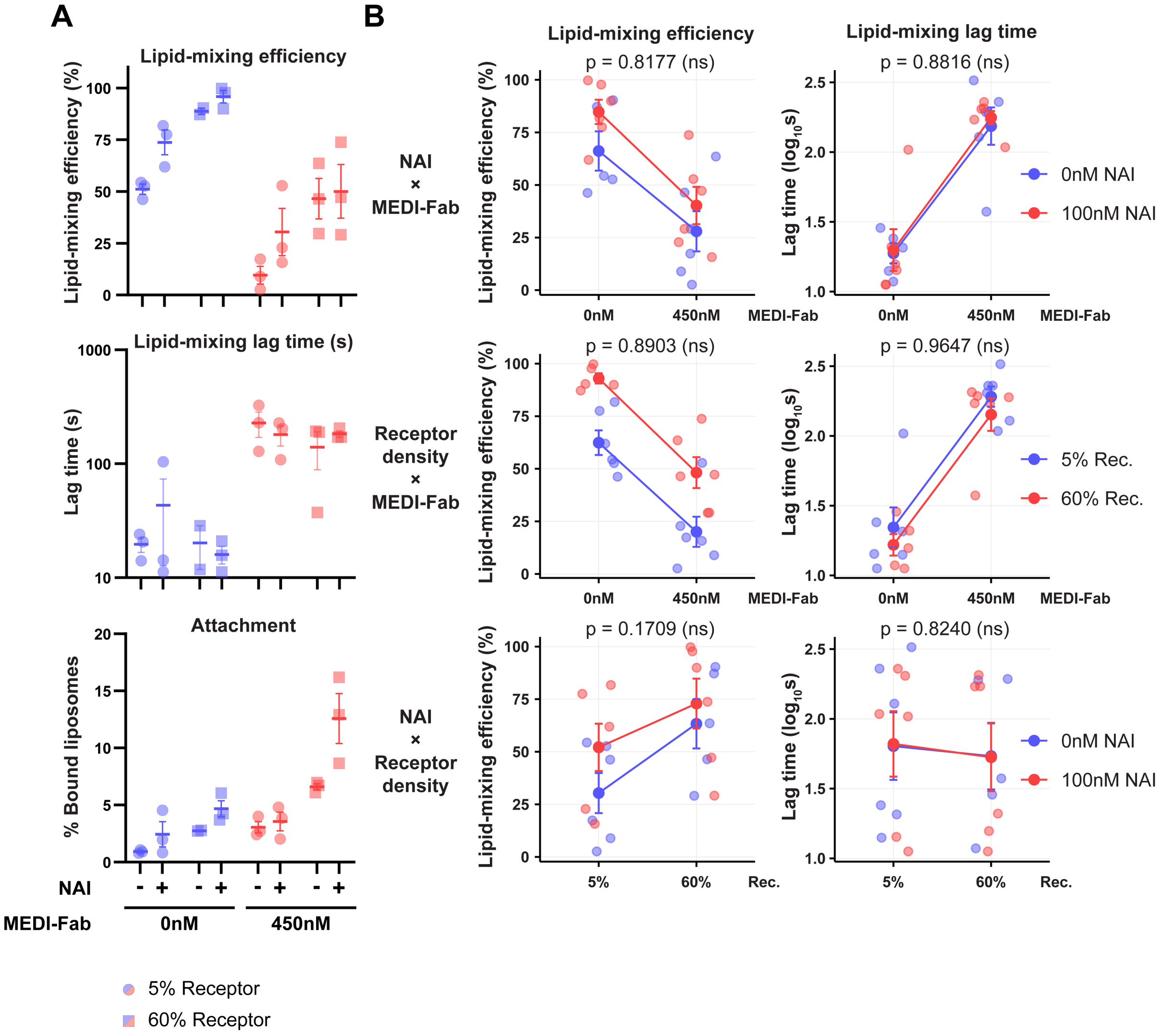


**Fig. S15. Inhibition of lipid-mixing by MEDI-Fab is insensitive to receptor context.** (A) Per-condition measurements of lipid-mixing efficiency, lag time, and attachment from the experiment in Fig 5C. (B) Experiment-wide interaction effects of NAI and MEDI (top), receptor density and MEDI (middle), and NAI and receptor density (bottom) on lipid-mixing efficiency (left column) and lag time (right column). Individual data points from three independent experiments were plotted with mean +/- SEM (Lipid-mixing efficiency and Attachment) or geometric mean +/- SEM (Lipid-mixing lag time). P-values for the interaction effects are derived from three-way ANOVA (ns: not significant) (see Supplementary Table 2). Lipid-mixing efficiency interaction plots for NAI x MEDI-Fab and Receptor Density x MEDI-Fab were replotted from Fig. 5C.

### Supplementary Tables

#### Supplementary Table 1: ssDNA Conjugates

| Receptor-ssDNA | | M.W. (g/mol) | Relevant Figures |
|---|---|---|---|
| 6’SL-DNA12 (2A’) | /5 3’SL/5’- TTT CCG TGT CGA -3’ | 4445.7 | S6, S9 |
| 6’SL-DNA24 (2A’-3A’) | /5 6’SL/5’- TCA ACA TCA ACA TTT CCG TGT CGA -3’ | 8086.5 | 3*E*, 4, 5, 5*B*, S7, S8, S9, S10, S12-15 Movie 2 |
| 3’SL-DNA12 (2A’) | /5 6’SL/5’- TTT CCG TGT CGA -3’ | 4445.7 | N/A |
| 3’SL-DNA24 (2A’-3A’) | /5 3’SL/5’- TCA ACA TCA ACA TTT CCG TGT CGA -3’ | 8086.5 | 4*B*, 4*C*, S12-13 |
| 6’SL-DNA14XL (1A’) | /5 6’SL/5’- CT CAG TGG ACA GCC -3’ | 5067.1 | N/A |
| 3’SL-DNA14XL (1A’) | /5 3’SL/5’- CT CAG TGG ACA GCC -3’ | 5067.1 | N/A |
| **Receptor-free ssDNA** | | **M.W. (g/mol)** | **Relevant Figures** |
| 12nt (2A’) | 5’- TTT CCG TGT CGA -3’ | 3616.31 | S6 |
| 18nt (2A’-3B’) | 5’- TCA ACA TTT CCG TGT CGA -3’ | 5432.44 | N/A |
| 24nt (2A’-3A’) | 5’- TCA ACA TCA ACA TTT CCG TGT CGA-3’ | 7248.57 | 3*E*, 4, 5, S7, S8, S10, S12-15 |
| **Lipid-ssDNA** | | **M.W. (g/mol)** | **Relevant Figures** |
| Lipid-DNA18 (2A-3B) | /5Lipid/5’- TCG ACA CGG AAA TGT TGA -3’ | 6206.7 | S6 |
| Lipid-DNA24 (2A-3A) | /5Lipid/5’- TCG ACA CGG AAA TGT TGA TGT TGA -3’ | 8140.9 | 3*E*, 4, 5, S7, S8, S9, S10, S12-15, Movie 2 |
| Lipid-DNA12XL (2A-2B) | /5Lipid/5’- TCG ACA CGG AAA AAAAA -3’ | 5884.5 | N/A |
| Lipid-DNA14XL (1A-1B) | /5Lipid/5’- GGC TGT CCA CTG AG TTTTT -3’ | 6455.8 | N/A |

**Oligonucleotide sequences and molecular weights in g/mol for receptor ssDNA, receptor-free ssDNA, and lipid ssDNA.** Mutually complementary sequences have the same color. For example, in the left column, 2A red sequences are complementary to 2A’ red sequences. Longer sequences might include subsequences complementary to more than one shorter sequence, shown with more than one color.

#### Supplementary Table 2: Three-way ANOVA of NAI vs. Receptor Density vs. MEDI-Fab

| Source of Variation | LM Efficiency | | LM Lag Time | | Binding | |
|---|---|---|---|---|---|---|
| **NAI** | 0.0324 | * | 0.7214 | ns | 0.0037 | ** |
| **Receptor Density** | 0.0001 | *** | 0.317 | ns | <0.0001 | **** |
| **MEDI-Fab** | <0.0001 | **** | <0.0001 | **** | 0.0001 | *** |
| **NAI x Receptor Density** | 0.1709 | ns | 0.824 | ns | 0.0615 | ns |
| **NAI x MEDI-Fab** | 0.8177 | ns | 0.8816 | ns | 0.3091 | ns |
| **Receptor Density x MEDI-Fab** | 0.8903 | ns | 0.9647 | ns | 0.0104 | * |
| **NAI x Receptor Density x MEDI-Fab** | 0.9411 | ns | 0.2991 | ns | 0.1017 | ns |

**Three-way ANOVA p-values and significance indicators for lipid-mixing efficiency, lipid-mixing lag time, and pre-pH-drop binding for the experiment presented in Fig 5*C*.** "x" indicates interaction between adjacent independent variables. ns $P > 0.05$, *$P < 0.05$, **$P < 0.01$, ***$P < 0.001$, ****$P < 0.0001$.

### Supplementary Datasets

#### Dataset 1: Sequencing of H36-26 Hybridoma Variable Region

BioIntron Biologic Inc. was contracted to sequence a hybridoma producing the antibody H36-26 IgG (Sb epitope). Total RNA was reverse transcribed into cDNA using anti-sense primers following the technical manual of Hiscript III Reverse Transcriptase (Vazyme, Cat# R302-01). Then the antibody fragments of VH and VL were amplified according to the SOP of Biointron Biologic Inc. The PCR fragments were cloned into TA/Blunt-Zero Cloning vector. Colony PCR was performed to screen for clones and no less than three positive clones were sequenced.

**1. Heavy Chain** (Leader sequence-FR1+CDR1+FR2+CDR2+FR3+CDR3+FR4)
**1) DNA Sequence: 414 bp**
ATGGCTTGGGTGTGGACCTTGCTATTCCTGATGGCAGCTGCCCAAATTATCCAGACA
CAGCTCCAGTTGGTGCAGTCTGGACCTGACCTGAAGAAGCCTGGAGAGACAGTCAGGATCTCCTGCAA
GGCTTCTGGTTATACCTTCACAGACTTTACAATGCACTGGGTGAAGCAGACTCCAGGAAAGGGTTTAAA
GTGGATGGGCTACATAAACACTGAGACTGGTGCGCCAACATATGCAGATGACTTCAAGGGACGGTTTG
CCTTCTCTTTGGAGACCTCTGCCAGCACTGCCTATTTGCAGATCAACAACCTCAAAGATGAGGACACGG
CTACATATTTCTGTACTAGAACGTATTATAGATTCACCTGGTTGGCTTACTGGGGCCAAGGGACTCTGG
TCTCTGTCTCTGCA

**2）Amino Acid Sequence: 138 aa**
MAWVWTLLFLMAAAQIIQT
QLQLVQSGPDLKKPGETVRISCKASGYTFTDFTMHWVKQTPGKGLKWMGYINTETGAPTYADDFKGRFAF
SLETSASTAYLQINNLKDEDTATYFCTRTYYRFTWLAYWGQGTLVSVSA

**2. Light Chain** (Leader sequence-FR1+CDR1+FR2+CDR2+FR3+CDR3+FR4 )
**1) DNA Sequence: 399 bp**
ATGGAATCACAGACTCAGGTCCTCATCTCCTTGCTGTTCTGGGTATCTGGTACCTGTGGG
GACATTGTGATGACACAGTCTCCATCCTCCCTGAGTGTGTCAGCAGGAGAGAAGGTCACTATGAACTG
CAAGTCCAGTCAGAGTCTGTTAAACAGTGGAAATCAAAAGAACTATTTGGCCTGGTACCAGCAGAAACC
AGGGCAGCCTCCTAAACTTTTGATCTACGGGGCATCCACTAGGCAATCTGGGGTCCCTGATCGCTTCA
CAGGCAGTGGTTCTGGAACCGATTTCACTCTTACCATCAGCAGTGTGCAGGCTGAAGACCTGGCAGTT
TATTACTGTCAGAATGATCATAATTATCCGCTCACATTCGGTGCTGGGACTAAGCTGGAGCTGAAA

**2)Amino Acid Sequence: 133 aa**
MESQTQVLISLLFWVSGTCG
DIVMTQSPSSLSVSAGEKVTMNCKSSQSLLNSGNQKNYLAWYQQKPGQPPKLLIYGASTRQSGVPDRFTG
SGSGTDFTLTISSVQAEDLAVYYCQNDHNYPLTFGAGTKLELK

#### Dataset 2: H36-26 (human IgG1) Sequence

BioIntron Biologic Inc. was contracted to clone the H36-26 variable regions (sequenced from a hybridoma, see SI Dataset 1) into a human IgG1 backbone. See the below structure map and corresponding sequences.

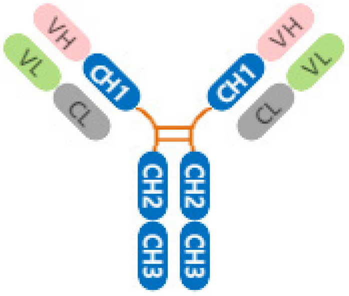


**Heavy chain sequences:** Signal peptide**-VH-**CH (Human IgG1 E356D M358L)
H36-26-hIgG1-HC
**GCGGCCGC**AAACTACAAGACAGACTTGCAAAAGAAGGC**ATG**CACAGCTCAGCACTGCTCTGTTGCCTG
GTCCTCCTGACTGGGGTGAGGGCC
**CGGCGAGACCGTGAGGATCTCCTGCAAGGCCTCCGGCTACACCTTCACCGACTTCACCATGCACTGG**
**GTGAAGCAGACCCCCGGCAAGGGCCTGAAGTGGATGGGCTACATCAACACCGAGACCGGCGCCCCC**
**ACCTACGCCGACGACTTCAAGGGCAGGTTCGCCTTCTCCCTGGAGACCTCCGCCTCCACCGCCTACC**
**TGCAGATCAACAACCTGAAGGACGAGGACACCGCCACCTACTTCTGCACCAGGACCTACTACAGGTT**
**CACCTGGCTGGCCTACTGGGGCCAGGGCACCCTGGTGTCCGTGTCCGCC**
ATCGGTCTTCCCCCTGGCACCCTCCTCCAAGAGCACCTCTGGGGGCACAGCGGCCCTGGGCTGCCTG
GTCAAGGACTACTTCCCCGAACCGGTGACGGTGTCGTGGAACTCAGGCGCCCTGACCAGCGGCGTGC
ACACCTTCCCGGCTGTCCTACAGTCCTCAGGACTCTACTCCCTCAGCAGCGTGGTGACCGTGCCCTCC
AGCAGCTTGGGCACCCAGACCTACATCTGCAACGTGAATCACAAGCCCAGCAACACCAAGGTGGACAA
GAAAGTTGAGCCCAAATCTTGTGACAAAACTCACACATGCCCACCGTGCCCAGCACCTGAACTCCTGG
GGGGACCGTCAGTCTTCCTCTTCCCCCCAAAACCCAAGGACACCCTCATGATCTCCCGGACCCCCGAG
GTCACATGCGTGGTGGTGGACGTGAGCCACGAAGACCCTGAGGTCAAGTTCAACTGGTACGTGGACG
GCGTGGAGGTGCATAATGCCAAGACAAAGCCGCGGGAGGAGCAGTACAACAGCACGTACCGTGTGGT
CAGCGTCCTCACCGTCCTGCACCAGGACTGGCTGAATGGCAAGGAGTACAAGTGCAAGGTCTCCAACA
AAGCCCTCCCAGCCCCCATCGAGAAAACCATCTCCAAAGCCAAAGGGCAGCCCCGAGAACCACAGGTG
TACACCCTGCCCCCATCCCGGGACGAGCTGACCAAGAACCAGGTCAGCCTGACCTGCCTGGTCAAAG
GCTTCTATCCCAGCGACATCGCCGTGGAGTGGGAGAGCAATGGGCAGCCGGAGAACAACTACAAGACC
ACGCCTCCCGTGCTGGACTCCGACGGCTCCTTCTTCCTCTACAGCAAGCTCACCGTGGACAAGAGCAG
GTGGCAGCAGGGGAACGTCTTCTCATGCTCCGTGATGCATGAGGCTCTGCACAACCACTACACGCAGA
AGAGCCTCTCCCTGTCTCCGGGTAAA**TGA**T**TCTAGA**
MHSSALLCCLVLLTGVRA
QLQLVQSGPDLKKPGETVRISCKASGYTFTDFTMHWVKQTPGKGLKWMGYINTETGAPTYADDFKGRFAFS
LETSASTAYLQINNLKDEDTATYFCTRTYYRFTWLAYWGQGTLVSVSA
ASTKGPSVFPLAPSSKSTSGGTAALGCLVKDYFPEPVTVSWNSGALTSGVHTFPAVLQSSGLYSLSSVVTVP
SSSLGTQTYICNVNHKPSNTKVDKKVEPKSCDKTHTCPPCPAPELLGGPSVFLFPPKPKDTLMISRTPEVTC
VVVDVSHEDPEVKFNWYVDGVEVHNAKTKPREEQYNSTYRVVSVLTVLHQDWLNGKEYKCKVSNKALPAPI
EKTISKAKGQPREPQVYTLPPSRDELTKNQVSLTCLVKGFYPSDIAVEWESNGQPENNYKTTPPVLDSDGSF
FLYSKLTVDKSRWQQGNVFSCSVMHEALHNHYTQKSLSLSPGK

**Light chain sequences:** Signal peptide **-VL-** CL (Human Kappa)
H36-26-hIgG1-LC
**GCGGCCGC**AAACTACAAGACAGACTTGCAAAAGAAGGC**ATG**CACAGCTCAGCACTGCTCTGTTGCCTG
GTCCTCCTGACTGGGGTGAGGGCC
**CTGGAGAGAAGGTGACCATGAACTGTAAGAGCTCCCAGTCCCTGCTGAACTCCGGCAACCAGAAGA**
**ACTACCTGGCCTGGTACCAGCAGAAACCCGGCCAGCCTCCTAAGCTGCTGATCTACGGCGCCTCTAC**
**CAGGCAGAGTGGCGTGCCTGATAGGTTCACCGGCTCCGGAAGCGGAACCGATTTCACCCTGACTATC**
**TCCTCCGTGCAGGCTGAGGACCTGGCAGTGTACTACTGCCAGAACGACCACAACTACCCCCTGACTT**
**TCGGCGCAGGCACCAAGCTGGAACTGAAG**
ATCTGATGAGCAGTTGAAATCTGGAACTGCCTCTGTTGTGTGCCTGCTGAATAACTTCTATCCCAGAGAG
GCCAAAGTACAGTGGAAGGTGGATAACGCCCTCCAATCGGGTAACTCCCAGGAGAGTGTCACAGAGCA
GGACAGCAAGGACAGCACCTACAGCCTCAGCAGCACCCTGACGCTGAGCAAAGCAGACTACGAGAAA
CACAAAGTCTACGCCTGCGAAGTCACCCATCAGGGCCTGAGTTCGCCCGTCACAAAGAGCTTCAACAG
GGGAGAGTGT**TGA**T**TCTAGA**
DIVMTQSPSSLSVSAGEKVTMNCKSSQSLLNSGNQKNYLAWYQQKPGQPPKLLIYGASTRQSGVPDRFTG
SGSGTDFTLTISSVQAEDLAVYYCQNDHNYPLTFGAGTKLELK
NNFYPREAKVQWKVDNALQSGNSQESVTEQDSKDSTYSLSSTLTLSKADYEKHKVYACEVTHQGLSSPVTK
SFNRGEC
